# Characterizing the diversity of L2/3 human neocortical neurons in epilepsy

**DOI:** 10.1101/2022.06.13.495678

**Authors:** J. Keenan Kushner, Paige B. Hoffman, Christine Brzezinski, Molly M. Huntsman, Allyson L. Alexander

## Abstract

In the current study, we performed whole-cell current clamp recordings from human cortical neurons in layer 2/3 of the human neocortex in order to characterize the diversity of L2/3 human neocortical neurons in epileptic foci with various etiologies in order to begin to elucidate the underlying mechanisms of hyperexcitability which are still mostly unknown. We differentiated neuronal subtypes based on their firing patterns and AHP kinetics or epilepsy subtype (malformation of cortical development (MCD) vs. other (non-MCD)). We found that L2/3 pyramidal neurons have diverse firing properties and action potential kinetics, with some neurons looking remarkably similar to LTS interneurons. We also saw that L2/3 pyramidal neurons could be split into those with fast AHPs and those without, medium AHPs (mAHPs). Based on these parameters, we were unable to significantly differentiate neurons based on firing properties indicating that AHP component kinetics alone do not dictate L2/3 pyramidal neuron firing in human epileptic cortical slices. We also report significant differences in intrinsic properties between MCD and non-MCD and control L2/3 pyramidal neurons and are the first to characterize that wash on of the proconvulsant drug, 4-aminopyridine (4-AP), leads to increased AP duration, less firing rate (FR) accommodation, and slowed down AHPs. Overall, the present study is the first to characterize the large variability of L2/3 human neocortical pyramidal neurons, to compare between L2/3 pyramidal neurons within the epileptic foci between MCD and non-MCD cases, to use control tissue from tumor patients without incidence of seizure, and to determine the influence of 4-AP on L2/3 pyramidal neuron intrinsic properties.

## Introduction

Most children with refractory epilepsy opt-in for resection which has been shown to provide long-term seizure control 40–80% of operative candidates (Krucoff et al., 2017)^1^. Refractory epilepsy or pharmacoresistant epilepsy is a debilitating disorder with a lexicon of environmental and genetic causes in addition to its refractory nature and our current inability to know the exact diagnosis until post biopsy processing and analysis of the cortical tissue. This makes it extremely hard to treat. However, researchers for the last four-five decades have been trying to understand the mechanisms and trademarks underlying each epileptic diagnosis in order to help these children before they need to go into a surgical suite. This work has included *ex vivo* electrophysiology, histology, and *in vivo* recording and imaging studies; however, the literature base is filled with many diverse types of experiments with little overlap of methods. We therefore set out to characterize L2/3 pyramidal neurons in slices from human epileptic foci resected in our local children’s hospital.

L2/3 is thought to control the gain of cortical output and modulates sensory evoked responses (Quiquempoix et al., 2018)^2^ making it a quintessential layer to explore for cellular and network abnormalities that may lead to seizure activity. Past research has looked into a wide variety of neurons across layer of different human cortical regions, however, most did not focus on a particular layer, others used cortical slices from outside the epileptic foci as “normal tissue”, and even some did some rudimentary characterization, but not enough to determine the characteristics of basic L2/3 neuron properties (Schwartzkroin et al., 1983, Avoli et al., 1989, Foehrig et al., 1991, Lorenzon et al., 1992, DeFelipe et al., 1992, Molnar et al., 2008, Mohan et al., 2015, Eyal et al., 2016, Beaulieu-Laroche et al., 2018, Kalmbach et al., 2018, Berg et al., 2020, Levinson et al., 2020, Moradi Chameh et al., 2021) ^3–15^. Therefore, we wanted to characterize basic intrinsic properties of L2/3 neocortical neurons from within the epileptic foci of human resected tissue.

In addition, there has been little to no research comparing neuronal properties between epilepsies. In this study, we wanted to determine the difference between malformation of cortical development epilepsies (MCDs epilepsy) including: focal cortical dysplasia (FCD) and Tuberous Sclerosis (TSC), which have been previously shown to have similar cell types and intrinsic properties (Cepeda et al., 2012, Cepeda et al., 2018)^16,17^, epilepsies that are not canonically cortical malformations (non-MCD) including: types of gliosis, polymicrogyria, low grade tumor’s that cause seizures, and temporal lobe epilepsies (TLE) and tumor control neurons that we receive from those going under resective surgery for tumor removal and have no history of seizures.

FCD is the main cause of medically intractable pediatric epilepsy and encompasses alterations of cortical neuron population percentages, the presence of dysmorphic neurons and balloon cells as well as abnormal cytoarchitecture. This has all been linked to initiation and development for seizure activity (Blumcke et al., 2011, Liang et al., 2020)^18,19^s. In addition to FCD and all of its subtypes, the MCD class of epilepsy also encompasses Tuberous Sclerosis (TSC), hemimegaloencephaly or schizencephaly (Curatolo et al., 2018)^20^. For most of the MCD epilepsies, especially FCD and TSC, the mammalian target of rapamycin (mTOR) pathway which drives the epileptogenesis which is triggers by genetic mutations along the pathway as well as other acquired factors (Cepeda et al., 2012, Curatolo et al., 2018, Wu et al., 2021, Heinemann et al., 2014, Jozwiak et al., 2006, O’Dell et al., 2012, Crino, 2015, Iffland and Crino, 2017, Lipton and Sahin, 2014, Park et al., 2018, Nagakawa et al., 2017, Guerrini et al., 2015)^16,20–30^. The non-MCD epilepsies don’t really contain vast subtypes of abnormal neuron types or dyslamination like the MCD epilepsies and instead manifest dysfunctional astrocytes and other glial cells that play crucial roles in uptake and redistributions of ions, neurotransmitters, and glucose, and therefore, help facilitate network communication and neuronal health and metabolism (Heuser et al., 2014)^31^. Based on the mechanistic, developmental, and cellular differences between MCD and non-MCD, we wanted to compare them against each other and against control tissue in order to determine if L2/3 pyramidal neurons differed in their intrinsic properties between epilepsy subtypes. We observed that L2/3 pyramidal neurons are similar between MCD and non-MCD and instead are quite different between the epilepsy subtypes and control L2/3 neurons.

Lastly, it is common to wash on 4-aminopyridine (4-AP), an A-type K+ channel blocker, to induce seizure like activity (ictal activity) in human cortical slices. Blocking with 4-AP (100μM) results in generation of a GABAergic, long-lasting depolarization (termed giant depolarizing potentials; Avoli and Perreault, 1987; Avoli et al., 1988)^32,33^, increase action potential (AP) duration (Williams and Hablitz, 2015)^34^, neurotransmitter release and elicits a large increase in extracellular [K+] and of course, interictal and ictal activity (Avoli et al., 2016)^35^. In addition, it has been shown to block fast AHP (fAHP) which is an AHP component also regulated by K+ currents and therefore, is blocked by 4-AP leading to reduced FR accommodation (Avoli et al., 1991, Lorenzon et al., 1992, Foehrig et al., 1991) ^8,10,36^. Overall, this study is the first to characterize L2/3 pyramidal neurons in depth including the differences in firing rate property, AHP kinetics, how they differ based on the epileptic subtype they belong to and is the first time a complete set of human L2/3 pyramidal neurons from epileptic foci intrinsic properties have been analyzed when 4-AP is washed on.

## Materials and Methods

### Acute human brain slice preparation for electrophysiology

Resected neocortical tissue was obtained from the local children’s hospital in collaboration with a network of neurosurgeons. All patients provided written informed consent to obtain neocortical tissue and demographic and surgical information collection. All information and experimental uses were in accordance with the respective hospital Institution Review Board before beginning the study. For the present study, cortical tissue samples were collected from June 2018 through May 2022. Cortical tissue samples from epilepsy patients of multiple etiologies were included; FCD type IA/B, FCD type IIA/B, FCD IIID, (Blumcke et al., 2009, Blumcke et al., 2011; Guerrini et al., 2015)^18,29,37^, MCD, TSC, and non-FCD etiologies including tumor, gliosis, polymicrogyria, and encephalomalacia. Patients underwent a variety of resection procedures in order to extract epileptic foci, or in the case of control, cortical tissue in order to gain access to pathology. Surgery was performed under local anesthesia. Electrocorticography (ECoG) was performed during epilepsy resection surgery for defining the location and extent of the epileptic focus.

Immediately following the surgical resection, resected cortical tissue was submerged in 0-4°C carbogenated (95%O_2_-5% CO_2_) N-methyl-D-glucamine (NMDG) substituted artificial cerebrospinal fluid (aCSF), adapted from Ting et al., 2018^38^, ((in mM): 92 NMDG, 2.5 KCl, 1.25 NaH_2_PO_4_, 30 NaHCO_3_, 20 4-(2-hydroxyethyl)-1-pip-erazineethanesulfonic acid (HEPES), 25 d-glucose, 2 thiourea, 5 Na-ascorbate, 3 Na-pyruvate, 0.5 CaCl_2_·4H_2_O and 10 MgSO_4_·7H_2_O, pH adjusted to 7.3-7.4 with concentrated hydrochloric acid (HCl), osmolality ^~^305mOsm/kg). The total duration from operating room to slicing was 15-20 minutes. The tissue was then placed in a carbogenated petri dish with 0-4°C carbogenated NMDG aCSF. Approximately 1 cm^3^ tissue blocks were sectioned, when applicable. Connective tissue and blood vessels were cleaned. No effort was made to remove the pia mater due to the risk of damaging the underlying grey matter. The ^~^1 cm^3^ tissue block was then superglued on to a vibratome stage (Leica Biosystems) and immersed in 0-4°C carbogenated NMDG aCSF. Acute coronal slices (400 μm) were prepared by slicing perpendicular to the pial surface to ensure dendrites of large pyramidal neurons were preserved. Slices were then incubated in carbogenated NMDG aCSF at ^~^35°C for 12 minutes and were then transferred to a room temperate (^~^23°C) carbogenated modified-HEPES aCSF ((in mM): 92 NaCl, 2.5 KCl, 1.2 NaH_2_PO_4_, 30 NaHCO_3_, 20 HEPES, 25 d-glucose, 2 thiourea, 5 Na-ascorbate, 3 Na-pyruvate, 2 CaCl_2_·4H_2_O, 2 MgSO_4_·7H_2_O, pH adjusted to 7.3-7.4 with HCl, osmolality ^~^305mOsm/kg). The slices remained in the ^~^23°C modified-HEPES aCSF for at least 30 minutes before being transferred to the recording chamber for electrophysiology experiments.

### Electrophysiology

For whole-cell patch clamp recordings, slices were placed in a submerged slice chamber and perfused at a rate of 2mL/min with heated (^~^33°C) recording aCSF ((in mM): 124 NaCl, 2.5 KCl, 1.2 NaH_2_PO_4_, 24 NaHCO_3_, 5 HEPES, 12.5 d-glucose, 2 CaCl_2_·4H_2_O, 2 MgSO_4_·7H_2_O, pH adjusted to 7.3-7.4 with HCl, osmolality ^~^305mOsm/kg). Slices were visualized using a moving stage microscope (Scientifica, SliceScope Pro 2000) equipped with X4 (0.10NA) and a X40 water immersion objective lens (0.80NA) objectives, differential interference contrast (DIC) optics, a SciCam Pro camera (Scientifica), and Micro-Manager 1.4 (Open Imaging). L2/3 pyramidal neurons were deciphered under DIC. Whole-cell patch clamp recordings were performed with 10cm fire polished borosilicate glass with filament pipettes (outer diameter: 1.5mm) (Sutter Instrument Cat# BF150-86-10) pulled to 3–6MΩ and filled with a potassium gluconate intracellular recording solution ((in mM): 135 potassium gluconate, 20 KCl, 10 HEPES,0.1 EGTA, 2 Mg-ATP, 0.3 Na_2_-GTP, pH adjusted to 7.3-7.4 with KOH, osmolality ^~^295mOsm/kg). Internal solutions also contained 3-5mg/mL (0.3%-0.5%) of biocytin (Sigma-Aldrich, CAS# 576-19-2) which was added the day of recording.

Electrical recordings were acquired with a MultiClamp 700B amplifier and were sent through a Hum Bug Noise Eliminator (Quest Scientific) to then be converted to a digital signal with the Axon^™^ Digidata^®^ 1440A digitizer using pCLAMP 10.7 software (Molecular Devices). Access resistance was monitored throughout the experiments and data were discarded if access resistance exceeded 25MΩ. No junction potential compensation or series resistance compensation was performed. In current clamp, compensation for voltage variations was achieved using a bridge balance circuit. Data were sampled at 10kHz.

### Electrophysiology experimental design

#### Ramped current injection

After achieving whole-cell configuration, L2/3 neurons were recorded from at rest in current clamp mode (I_hold_= 0pA). Following a three second baseline period, the holding current was linearly ramped from 0 to 1000pA over 2s. A total of 25 sweeps of data were collected for each neuron, and the data were used to determine neuron resting membrane potential (RMP), AP threshold, and rheobase current.

#### Square current injection

Following ramped current injections, we recorded the responses of L2/3 neurons to a series of square hyperpolarizing and depolarizing current injections. Before initiation of the series of current injections, the resting membrane potential of neurons was adjusted to approximately −60mV. Each cell was subjected to two series of 600ms square current injections: −100 to +100pA at 10pA intervals and −250-1000pA at 50pA intervals. Separate recordings of 1000ms square current steps from-250-1000pA at 50pA intervals was used to assess frequency adaptation over time. The data collected in these experiments were used to determine active and passive membrane properties of the neurons.

### Definitions of electrophysiologic parameters

#### Resting Membrane Potential

Resting Membrane Potential (RMP) was defined as the mean RMP (I_hold_= 0pA) during a 500ms baseline across all sweeps in the ramped injection experiments.

#### AP threshold

AP threshold was defined as the voltage at which dV/dt exceeded 20 V/s. AP threshold was calculated at the first AP of each sweep in the ramped injection experiments.

#### Rheobase current

Rheobase current was defined as the mean current injected at AP threshold for the first AP across all sweeps in the ramped injection experiments.

#### Input resistance (MΩ)

Input resistance was defined as the slope of the best fit line of the I-V plot using the −100 to +100pA series of current injections. Mean voltage response to each current injection step was defined as the difference between baseline mean membrane voltage (100ms before current injection) and the mean membrane voltage during the 100ms period from 50ms after the start of the injection to 150ms after the start of the current injection. This 100ms window was chosen to allow for measurement of the change in RMP after the membrane had charged and before any potential HCN channel activation. The IV plot was constructed using all current steps below rheobase.

#### Max firing rate

Max firing rate was defined as the number of action potentials (APs) over the length of the square current at the most depolarized current injection step before attenuation of firing rate. Max firing rate was calculated using the −250 to +1000pA series of current injections.

#### AP amplitude

Action potential amplitude was defined as the voltage difference between threshold (set at dV/dt = 20V/s) and AP peak. AP amplitude was calculated at the rheobase+100pA sweep of the −250 to +1000pA series of current injections.

#### AP halfwidth

Action potential halfwidth was defined as the time between the half-amplitude point on the upslope of the AP waveform to the half-amplitude point on the downslope of the AP waveform. AP halfwidth was calculated at the rheobase sweep of the −250 to +1000pA series of current injection.

#### After-hyperpolarization potential (AHP) magnitude

AHP magnitude was defined as the differences between the most hyperpolarized membrane voltage of the AHP (within 100ms after AP threshold) and AP threshold. AHP magnitude was calculated at both the rheobase sweep and the rheobase+100pA sweep of the −250 to +1000pA series of current injection.

#### AHP latency

AHP latency was defined as the time from AP threshold to the peak of the AHP. AHP latency was calculated at both the rheobase sweep and the rheobase sweep of the −250 to +1000pA series of current injection.

#### ΔAHP

ΔAHP was defined as the difference between the first and last AHP (ΔAHP = AHP_last_ – AHP_first_). ΔAHP was calculated at both the rheobase sweep and the rheobase+100pA sweep of the −250 to +1000pA series of current injection.

#### AP phase plot

The AP phase plot was obtained by plotting the rate of change of the mean AP for each cell from the rheobase sweep and the rheobase+100pA sweep of the −250 to +1000pA series of current injections as a function of the corresponding membrane voltage.

#### Firing rate accommodation ratio (FR ratio)

FR ratio was defined as the ratio of the first and the last ISIs, such that firing rate adaptation = ISI_first_/ISI_last ISI_. FR ratio was calculated at all current steps of the −250 to +1000pA series of current injections.

#### Instantaneous frequency plots

The initial and final instantaneous frequency plots were defined as 1/ISI_first_ or 1/ ISI_last_. Both instantaneous frequency plots were calculated at all current steps of the −250 to +1000pA series of current injections.

#### Frequency vs time plot

Frequency vs time plot was performed using the 1000ms −250 to +1000pA series of current steps. Frequency was defined as 1/ISI from each 100ms bin.

#### AP broadening ratio

AP broadening ratio was defined as the ratio of the AP halfwidths of the first two APs (broadening = halfwidth_second_/halfwidth_first_). The broadening ratio was calculated at both the rheobase sweep and the rheobase+100pA sweep of the −250 to +1000pA series of current injection.

#### AP amplitude adaptation ratio

AP amplitude adaptation was defined as the ratio of the AP amplitude of the average of the last three APs and the first AP, such that AP amplitude adaptation = mean amplitude_last 3 APs_/amplitude_first AP_. The AP amplitude adaptation ratio was calculated at both the rheobase sweep and the rheobase+100pA sweep of the −250 to +1000pA series of current injection.

#### Membrane decay τ (tau)

Membrane decay τ was determined by using a single exponential fit, f(t) = A*e*^-*t*/τ^to fit the change in RMP induced by a 100pA sweep in the −100 to +100pA (10pA steps) series of current injections.

#### Hyperpolarization induced sag (voltage sag ratio)

Hyperpolarization-induced sag was calculated using the equation, 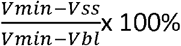, where V_min_ was defined as the most hyperpolarized membrane voltage during the current injection, V_ss_ was defined as the mean steady-state membrane voltage (last 200ms of the current injection), and V_bl_ was defined as the mean baseline membrane voltage (100ms before current injection). Hyperpolarization-induced sag was measured from the −250pA current injection.

### Extracellular application of 4-AP

In a subset of experiments, after the control ramp and current steps were completed, 100μm of 4-AP (Sigma-Aldrich, CAS#: 504-24-5) was washed on for 3 minutes (the amount of time it took to regularly induce 4-AP oscillations. After this time, the current step protocols were run again in order to compare intrinsic properties before and after 4-AP wash on.

### Histological methods

#### Biocytin filling neurons

All pipettes used for recording were filled with biocytin 3-5mg/mL (0.3%-0.5%). After current clamp recordings were complete, we slowly moving the recording pipette in small steps, alternating upward (along the Z-axis) and outward (along the X-axis) in voltage-clamp mode until establishing a gigaseal (GΩ), indicating successful re-sealing. After the gigaseal (GΩ) was obtained, the pipette was quickly maneuvered off of the cell. The slice was then left to rest in the recording chamber for 4-5 minutes to ensure transport of biocytin to distal dendrites and axon processes (Swietek et al., 2016)^39^ before the slice was moved into 4% PFA (Electron Microscopy Sciences, CAS #50-00-0) diluted in 0.1M Dulbecco’s Phosphate Buffered Saline (PBS) solution (Lonza, Cat# 12001-664). The slices stayed in the 4% PFA solution for 24-48 hours before being places in a 0.1M PBS solution until histology was performed.

#### Immunohistochemistry

Slices were stained within 3 weeks of recording day. First, slices were washed 3 times for 5 minutes each in 0.1M PBS. They were then incubated for 1 hour in blocking buffer (0.3% triton-X and 5% donkey serum diluted in 0.1M PBS). Slices were then washed again 3 times for 5 minutes each in 0.1M PBS before being incubated in blocking buffer with 1:1000 Alexa Fluor^®^ 594 Streptavidin (Jackson Immunoresearch) at 4°C overnight. The following day, slices were washed 3 more times for 5 minutes each in 0.1M PBS before being float mounted onto SuperFrost Plus microscope slides (Fisher) using a brush. Since 400μm slices are thick, 2 layers of electrical tape were used to create a square in which the slice sat in order to avoid smushing during imaging. Slices were then dried with a Kimwipe (Kimtech Science) and then Fluoromount-G^®^ was added before cover slipped (Fisher microscope cover glass).

#### 3D reconstruction of neuron morphology

Biocytin filled neurons were visualized using a Zeiss Axio Imager M2 microscope with excitation of 594nm and a 20X and 40X objective. 3D slide scanning was performed using Neurolucida (version 2021.1.3) under 40X magnification. Neuron tracing was performed using the Neurolucida neuron tracing software. Axons and dendrites were differentiated based on diameter and whether they had spines or boutons. The distance from pial surface was determined by the Neurolucida measuring tool. Layers were determined based on DIC imaging of the neuron shape and population density.

### Statistical analyses

All data analysis was performed using custom written MATLAB code, GraphPad Prism and Easy Electrophysiology v4.2.0 (http://www.easyelectrophysiology.com/). If data sets contained at least 10 data points, a D’Agostino & Pearson omnibus K2 normality test was performed to assess normality. For statistical tests between three or more groups with normal distribution, 10 or more data points, and equal variance, an ANOVA test was performed. Equal variance was determined using the Brown-Forsythe test. If data was normally distributed by had unequal variance (Brown-Forsythe test p<0.05, then a Welch’s ANOVA test was performed. Differences between groups was determined using a Tukey honestly significant difference (HSD) post hoc test. For statistical tests between three or more groups of non-normally distributed data, or data sets with < 10 data points, a Kruskal-Wallis test was performed. Differences between groups was determined using a Dunn’s multiple comparisons test and the *H*(degrees of freedom) statistic was checked against the critical chi-square value. For comparing intrinsic properties based on AHP shape (Figure 3), normal data with SD between groups < 2X were compared using an unpaired t-test, and non-normal data, and data with SD > 2X between groups, were compared using a Mann-Whitney U test. For comparing neuron intrinsic properties before and after 4-AP wash-on (Figure 5), a paired t-test was performed on normally distributed data with SD between groups being < 2X, and a Wilcoxon matched-pairs signed rank test was performed on non-normal data and data where SD between groups was > 2X.

**Figure 1:**
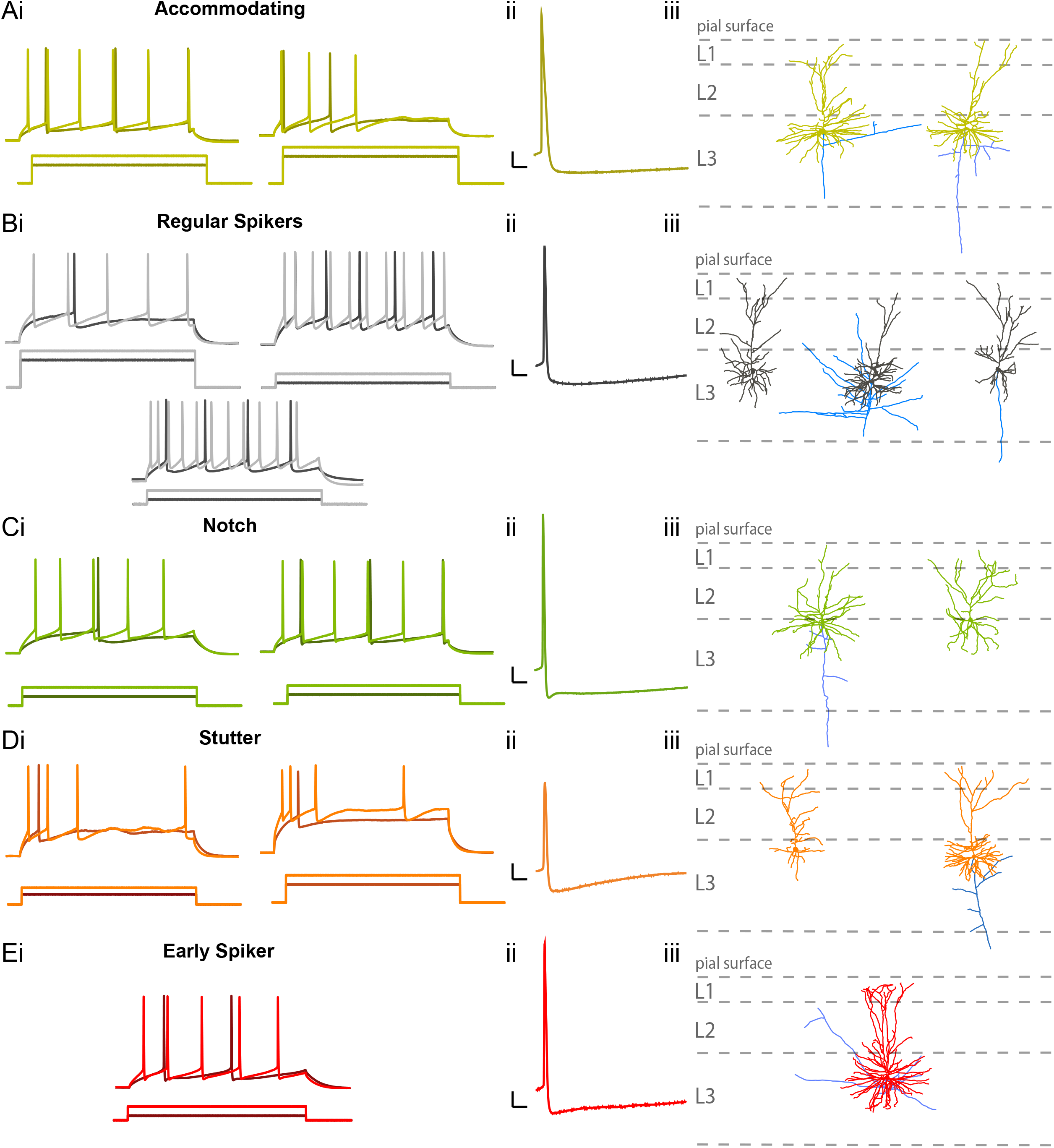
L2/3 neocortical neurons from human epileptic foci have diverse morphology and spiking properties. Ai-Ei) Depolarizing current steps at rheobase (dark trace) and rheobase +100 pA (light trace) elicit action potentials of various action potential shapes and frequency. Each group was determined based on firing property and/or action potential AHP shape. (Aii-Eii) Example of each action potential from neuronal subtype. Aiii-Eiii) Corresponding neuron traces with dendrites in neuronal subtype color scheme and axon branches in blue. Scale bars (10 mV, 10 msec).

**Figure 2:**
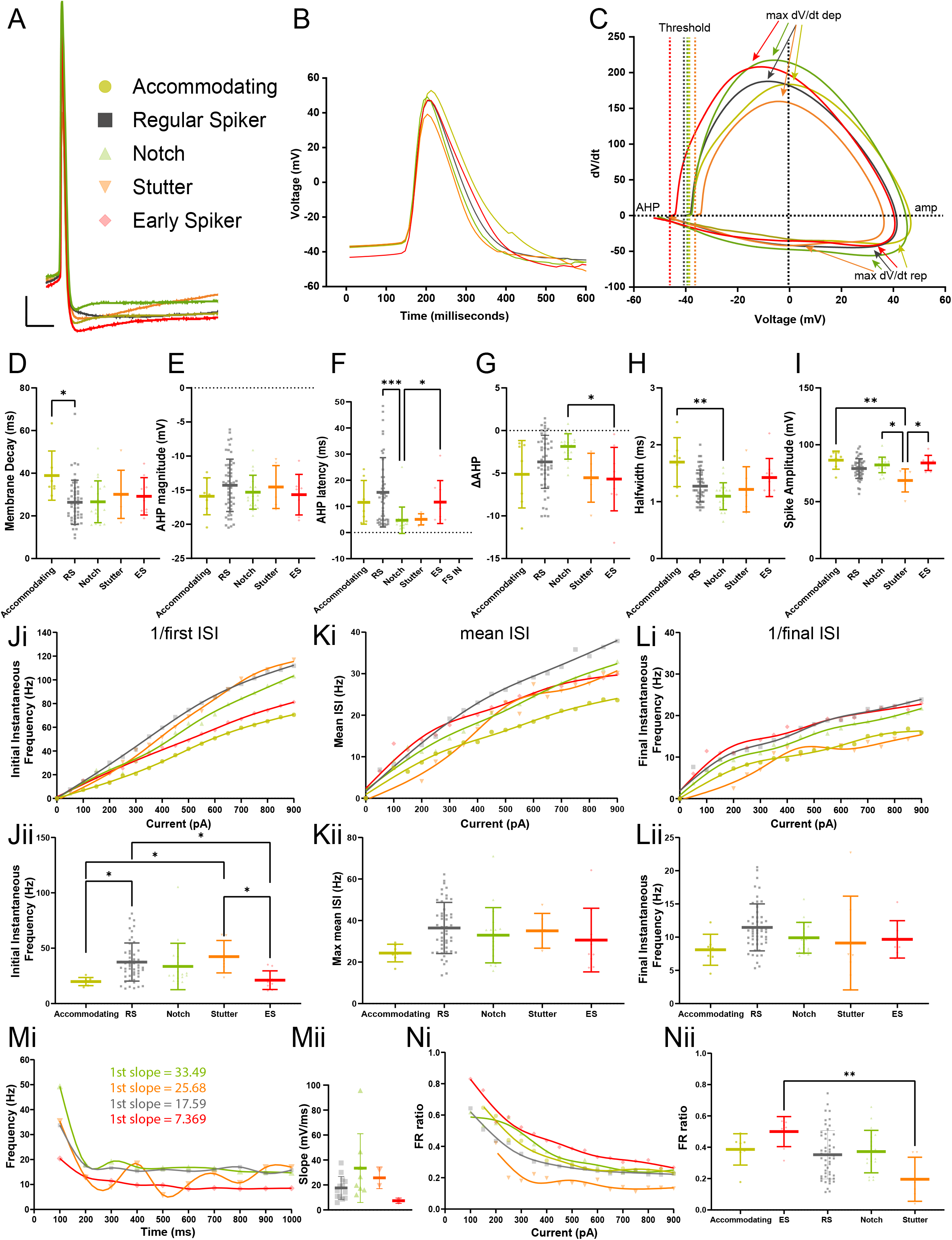
L2/3 putative pyramidal neuron subtypes show subtle differences in action potential kinetics and firing properties. A) Overlay of representative traces of L2/3 neuron subtype action potentials. Scale bar – 10 mV, 10ms. B) Overlay of average action potential of each neuron subtype. C.) Overlay of average phase plot dV/dT vs voltage (mV) indicating differences in subtype AP kinetics. D.) Membrane Decay (tau, ms) indicates difference between accommodating and RS neurons [KW test: p= 0.0468, H(4)= 9.650, Dunn test: accommodating vs RS (p= 0.0314, mean rank difference (MRD)= 31.49, z= 2.954)]. E.)After hyperpolarization potential (AHP) magnitude (mV) was similar between neuron subtypes. F.) Notch and stutter neurons have the shortest AHP latency (ms) with notch neurons having a significant shorter AHP latency compared to RS and ES neurons [KW test: p= 0.0002, H(4) =22.59, Dunn test: notch vs RS (p= 0.0001, MRD= 31.90, z= 4.409), notch vs ES (p= 0.0293, MRD= 32.49, z= 2.974)]. G.) Notch neurons have the most depolarized change in AHP magnitude (ΔAHP) over the rheobase +100 pA 600 ms depolarizing current step with a statistically more positive ΔAHP compared to ES neurons [KW test: p=0.0140, H(4) = 12.50, Dunn test: p= 0.0373, MRD= 31.33, z= 2.900]. H.) Notch neurons showed the shortest halfwidth (ms) with a significant difference compared to accommodating neurons [KW test: p=0.0052, H(4) = 14.76, Dunn test: p=0.0053, MRD= 40.79, z= 3.464]. I.) Stutter neurons showed a reduction in AP spike amplitude (mV) [KW test: p= 0.0029, H(4) = 16.11, Dunn test: stutter vs accommodating (p= 0.0052, MRD= 51.57, z= 3.472), stutter vs notch (p= 0.0317, MRD=3 6.89, z= 2.951), stutter vs ES (p= 0.0256, MRD= 42.44, z= 3.016)]. J.) Initial instantaneous firing frequency differed between subtypes (Hz) [KW test: p= 0.0005, H(4) = 19.86] (Table 2). K.) No post hoc differences in max mean firing rate (Hz) [KW test: p= 0.0422, H(4) = 9.896, no significant Dunn post hoc test]. L.) No post hoc differences in final instantaneous firing (Hz) [KW test: p= 0.0191, H(4) = 11.77, no significant Dunn post hoc test]. Ji-Li.) Initial, mean, and final frequency (Hz) plot vs injected current (pA). Jii-Lii.) Initial, mean, and final frequency (Hz) taken at the rheobase+100pA 600 ms current step. Mi-Mii.) Frequency (Hz) vs time (ms) plot indicating initial frequency differences (Hz) between subtypes indicated by slope (Hz/sec) above. Ni.) Firing Rate ratio vs injected current (pA) plot per neuron subtype. Nii.) Firing rate ratio of each neuron subtype taken at rheobase+100 pA indicates a lack of firing rate accommodation of ES neurons which was significantly less so than the FR ratio of stutter neurons. [KW test: p= 0.0054, H(4) = 14.69, Dunn test: (stutter vs ES neurons, p= 0.0021, MRD= −52.88, z= 3.707)].

**Figure 3:**
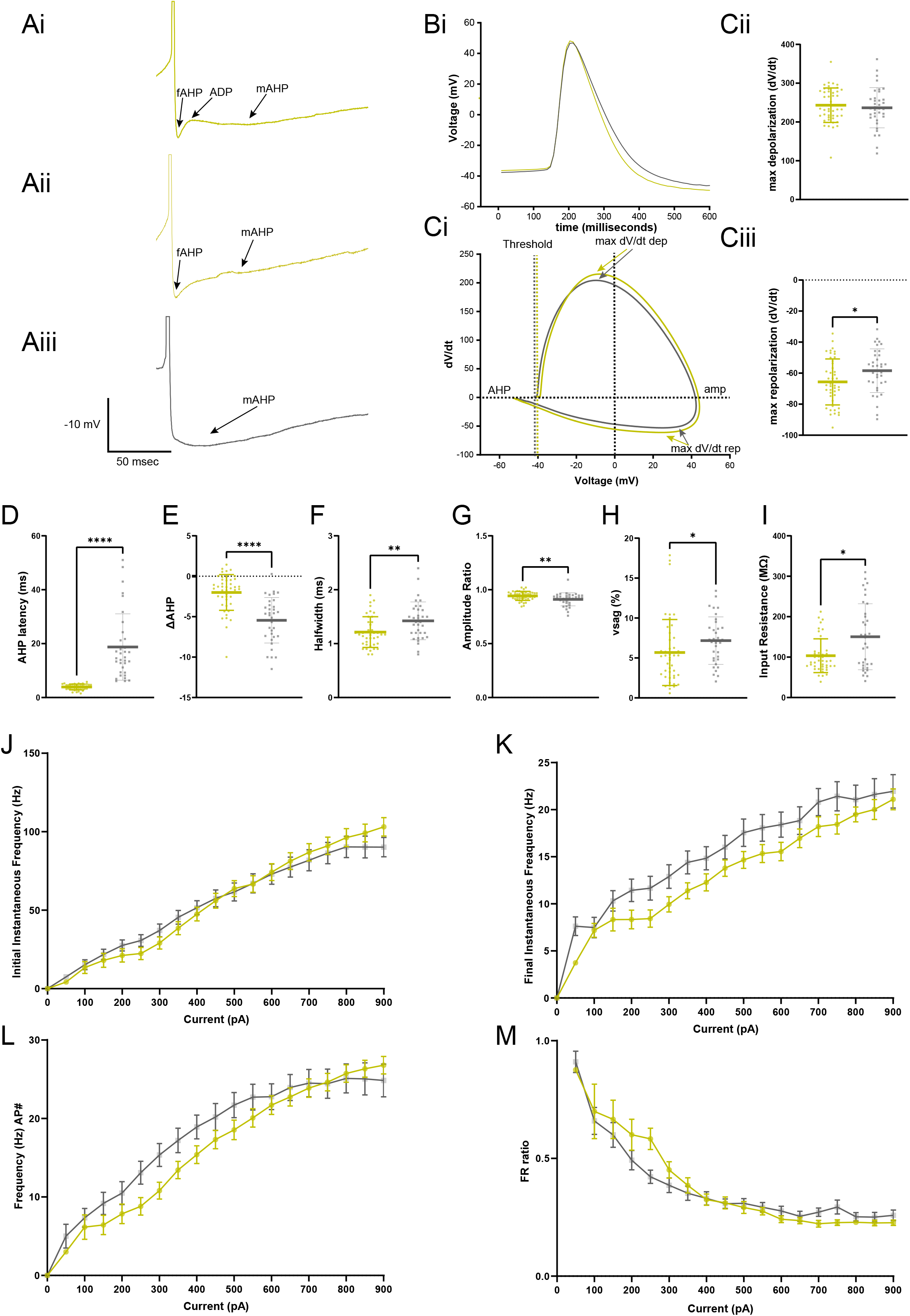
L2/3 pyramidal neurons with fast AHP or medium AHP do not differ in their firing rates or firing rate accommodation. Ai-Aiii) Example traces of L2/3 pyramidal neuron APs with Ai.)fAHP followed by ADP and mAHP, Aii.) fAHP only followed by mAHP and Aiii.) mAHP only. Bi.) Average action potential of fAHP and mAHP neuron subtypes. Ci.) Average phase plots. Cii.) Max depolarizing slope (dV/dT) for each neuron subtype indicates no significant differences between voltage-gates Na+ channels. Ciii.) Max repolarization slope (dV/dT) for each subtype indicates an increase in fAHP neuron repolarization kinetics which is related to voltage-gated K+ channels (unpaired t-test: p= 0.030, mean difference (MD)= 7.236 dV/dT, 95%CI= 0.7361 to 13.74). D.) AHP latency (ms) is shorter for fAHP neurons as expected (MWU test: p < 0.000001, MD= 10.59, 95% confidence interval(CI)= 9.050 to 13.60). E.) More depolarized ΔAHP of fAHP neurons (MWU test: p <0.0001, MD= −3.204, 95%CI= −4.608 to −2.289). F.) Shorter AP halfwidth (ms) halfwidths (unpaired t-test: p= 0.006, MD= 0.208ms, 95%CI= 0.0615 to 0.354). G.) Enhances fAHP neuron AP amplitude adaptation (unpaired t-test: p= = 0.006, MD: − 0.0322, 95%CI= −0.055 to −0.009). H.)Reduction in fAHP neuron voltage sag (%) which is related to HCN1 channel function and Ih current (MWU: p= 0.0115, MD: 1.922, 95%CI= 0.5720 to 3.405). I.) Lower input resistance of fAHP neurons perhaps due to more leak channels compared to mAHP neurons (MWU: p= 0.0323, MD= 50.17, 95%CI= 2.587 to 76.10). J.) No differences in initial instantaneous firing (Hz) ± SEM vs current (pA) steps, K.) maximum mean firing rate (Hz) ± SEM vs current (pA) steps, L.) final instantaneous firing (Hz) ± SEM vs current (pA) steps or M.) FR ratio between fAHP and mAHP neurons ± SEM vs current (pA) steps (Table 3).

**Figure 4:**
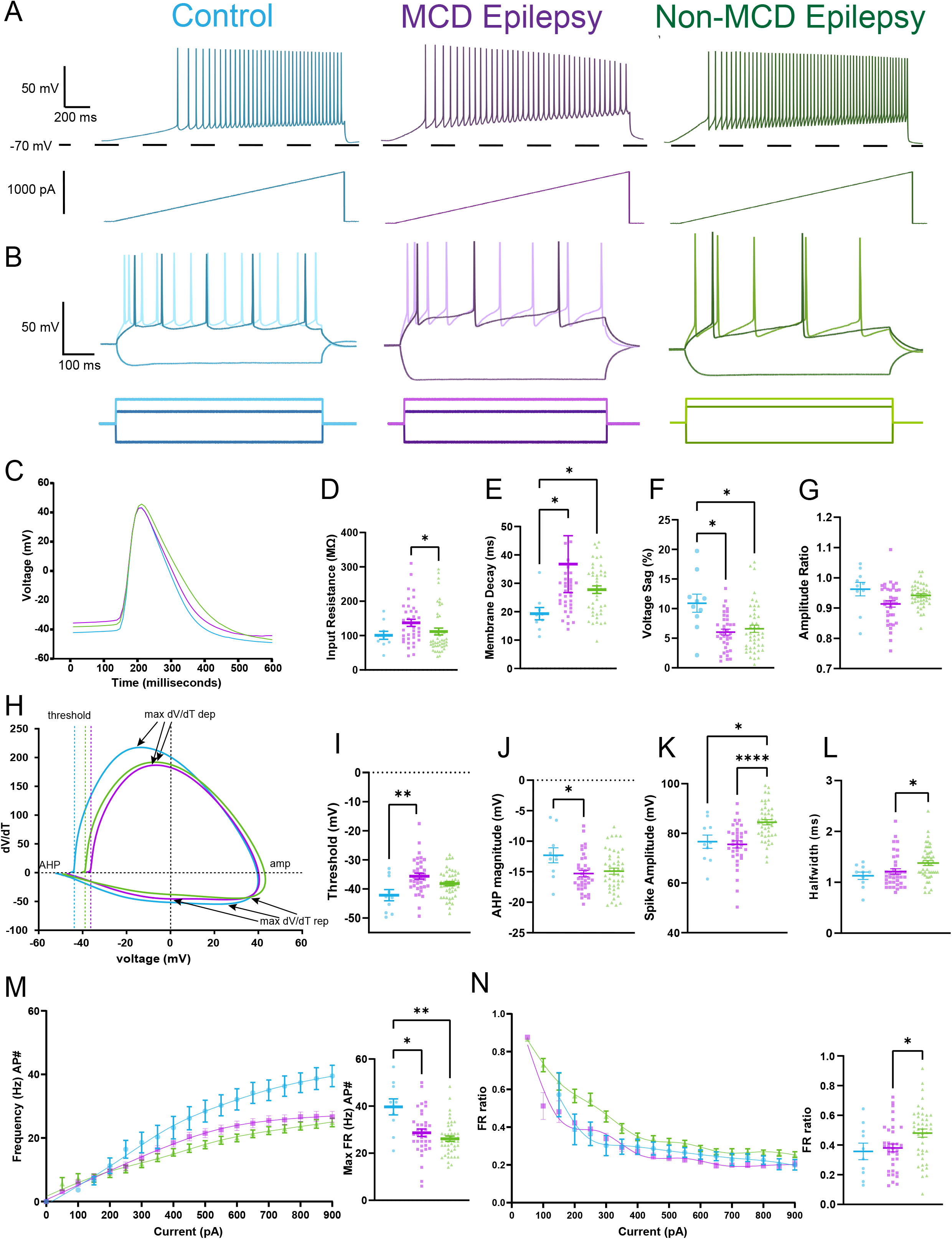
L2/3 epileptic pyramidal neurons show minor differences between each other, but crucial differences compared to control neurons. A) Example traces from current (pA) ramp protocol (1000 pA, 1 sec) of L2/3 pyramidal neurons from control (blue), MCD epileptic (purple) and non-MCD epileptic tissue (green). B) Example traces from L2/3 pyramidal neurons from control and epileptic subtype in vitro slices showing the 600 ms −250 pA hyperpolarizing step (darker negative trace) and 600 ms rheobase (pA) (darker positive trace) and rheobase+100 pA (lighter positive trace) depolarizing current steps (pA,600 msec steps) performed. C.) Average action potential of control and epileptic subtype L2/3 pyramidal neurons. D.) No differences in input resistance between control and epileptic subtype [KW test: p= 0.0287, H(2)= 7.101, Dunn test: (control vs MCD, p= 0.5066, MRD= −13.24, z= 1.376), (control vs non-MCD, p> 0.9999, MRD= 2.357, z= 0.2502)]. Only difference between MCD and non-MCD neurons (MCD vs non-MCD, p= 0.0267, MRD= 15.59, z= 2.616). E.) Elevation of membrane decay of L2/3 epileptic pyramidal neurons [KW test: p= = 0.0216, H(2) = 7.674, Dunn test: (control vs MCD, p= 0.0059, MRD= −22.79, z= 2.407), (control vs non-MCD, p= 0.0177, MRD= −25.45, z= 2.754)]. F.) Decrease in voltage sag of L2/3 epileptic pyramidal neurons [KW test: p= 0.0111, H(2)= 8.994, Dunn test: (control vs MCD, p= 0.0115, MRD= 27.82, z= 2.892), (control vs non-MCD, p= 0.0154, MRD= 26.35, z= 2.798)]. G.) No difference in AP amplitude adaptation ratio [W (DFn, DFd)= 3.316 (2.000, 22.14), p = 0.0550]. H) Average phase plots. I.) L2/3 MCD pyramidal neurons have depolarized AP threshold (mV) compared to control [ANOVA: p= 0.0032, F= 6.139, Tukey HSD: control vs MCD, p= 0.0033, MD= −6.520mV, 95%CI= −11.15 to −1.889)] J.) L2/3 MCD pyramidal neurons have increased AHP magnitude (mV) [ANOVA: p= 0.0434, F= 3.250, Tukey HSD: (control vs MCD, p= 0.0350, MD= 2.985mV, 95%CI= 0.1702 to 5.800)]. K.)L2/3 non-MCD pyramidal neurons have increased AP amplitude [KW test: p< 0.0001, H(2) = 22.71, Dunn test: (control vs non-MCD, p= 0.0287, MRD= −24.14, z= 2.591), MCD vs non-MCD, p <0.0001, MRD= −27.13, z= 4.566)] (mV) compared to MCD and control neurons. L) Significant longer AP halfwidths (ms) for L2/3 pyramidal neurons in the non-MCD subtype compared to MCD L2/3 pyramidal neurons [KW test: p= 0.0082, H(2) = 9.615, Dunn test: (MCD vs non-MCD, p= 0.0130, MRD= −16.97, z= 2.854)], with trends toward longer APs in epilepsy neurons compared to control. M., left) Mean ISI frequency (Hz) ± SEM vs injected current (pA). M., right) Max firing rate (Hz) taken at 900 pA depolarizing current step indicates reduction in max firing rate for both epileptic subtypes compared to control ([KW: p= = 0.0017, H(2) = 12.79, Dunn test: (control vs MCD, p= 0.0327, MRD= 23.68, z= 2.546), (control vs non-MCD, p= 0.0012, MRD= 32.47, z= 3.544)]. N., left) Firing Rate ratio ± SEM vs injected current (pA). N. right) Firing rate ratio taken at rheobase+100 pA indicates a significant lack of FR accommodation by non-MCD L2/3 neurons compared to MCD neurons [ANOVA, F = 4.223, p =0.0178,Tukey HSD: ( p= 0.0319, MD= −0.09999, 95%CI= −0.1929 to −0.007073) with no differences between epilepsy subtypes compared to control (Tukey HSD: (control vs MCD, p= 0.9244, MD= −0.02319, 95%CI= −0.1695 to 0.1232), (control vs non-MCD, p= 0.1065, MD= −0.1232, 95%CI= −0.2665 to 0.02011)).

**Figure 5:**
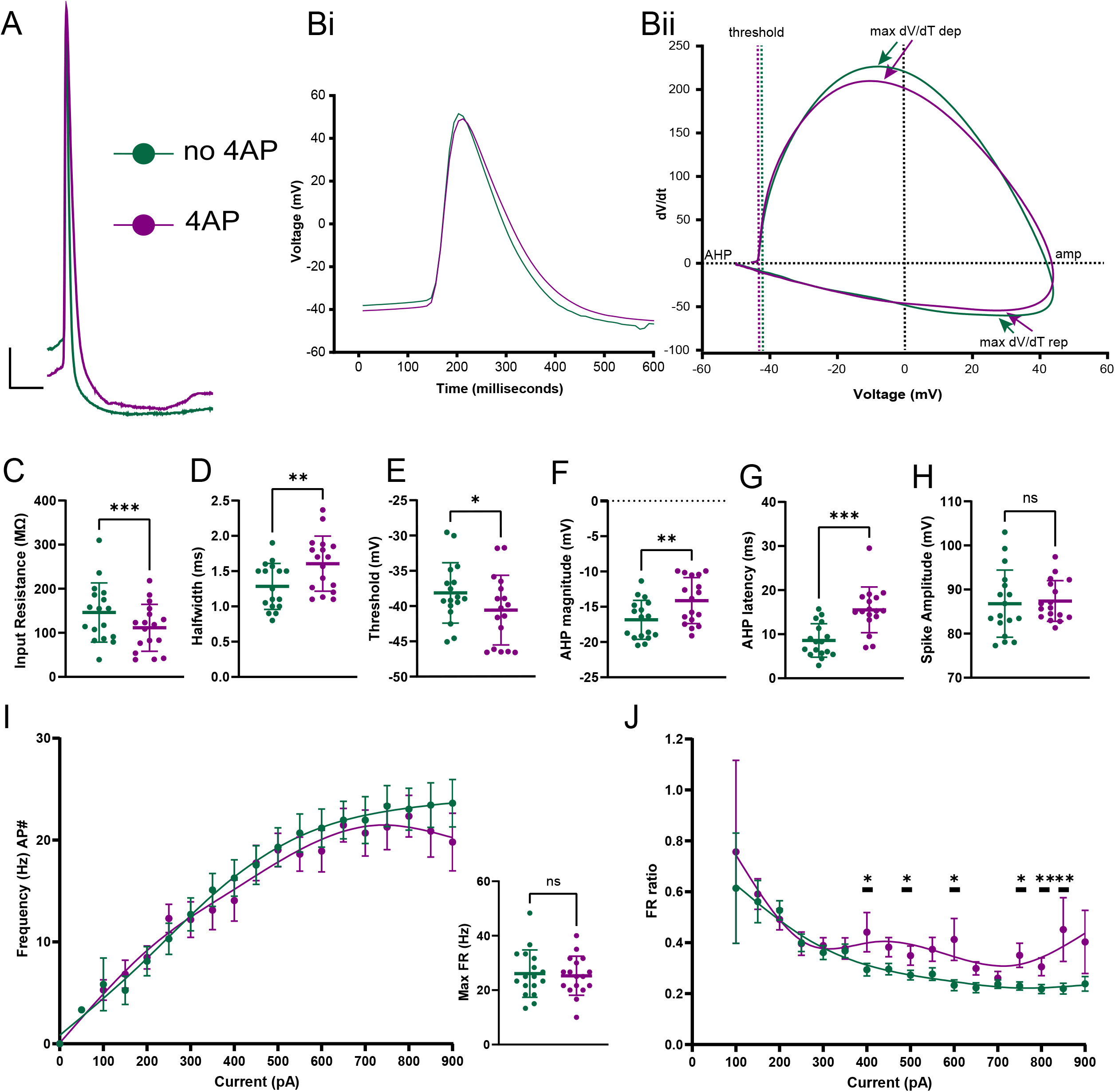
4-AP increases L2/3 pyramidal neuron AP AHP latency and AP halfwidth and decreased AHP magnitude leading to sustained firing. A.) Overlay of example AP before and after 4-AP wash on. Scale bar-10 mV, 10ms. Bi.) Overlay of average action potential before and after 4-AP wash on. Bii.) Overlay of phase plots. C.) Decrease in input resistance after 4-AP wash on (paired t-test: p= 0.0002, MD= −34.64ms, 95%CI= −50.10 to −19.19 MΩ). D.) Increase in AP halfwidth after 4AP wash on (paired t-test: p= 0.0091, MD= 0.3227, 95%CI= 0.09211 to 0.5534). E.) More hyperpolarized AP threshold (mV) after 4AP wash on (paired t-test: p= 0.0433, MD=-2.450mV, 95%CI= −4.816 to −0.08288). F.) Reduction of AHP magnitude after 4 AP wash on (Wilcoxon test: p= 0.0021, median of difference= 2.164, 95%CI= 1.122 to 3.682). G.) Longer AHP latency after 4AP wash on (paired t-test: p= 0.0002, MD= 6.940ms, 95%CI= 3.846 to 10.03). H.) No difference in AP amplitude (mV) (paired t-test: p= 0.7729). I.) No difference in max FR (Hz) ± SEM vs injected current (pA) (paired t-test: p= 0.3543). J.) Less FR accommodation± SEM vs injected current (pA) after 4AP wash on at 400, 500, 600, 750, 800, and 850 pA depolarizing current steps (see results for significance).

All statistical tests were two-tailed. Unless otherwise stated, experimental numbers are reported as *n*⍰=⍰x, y, where x is the number of neurons and y is the number of patients. Data visualizations were created in MATLAB R2018a, GraphPad Prism 9.3.1, Adobe Illustrator and OriginPro 2022 (Origin Lab). Data are presented as the mean ± SD unless otherwise stated as standard error of the mean (SEM). Frequency and FR ratio line graphs indicate smoothed data using the smoothing spline method in GraphPad Prism. Phase plots were created using the B-spline connect interpolation in OriginPro 2022.

## Results

### Cohort

Whole-cell current clamp recordings were obtained from human neocortical neurons located in L2/3 of acute brain slices collected from 28 epilepsy and 2 tumor control patients who underwent resective surgery. Patients had an average age of 10.3 years old and a range from 1–21 years old. Of these 30 patients, 17 were male and 13 were female. The brain samples analyzed in the present experiments were from the temporal (n = 15), frontal (n = 13), parietal (n = 3), or occipital (n = 1) neocortex. Anonymized patient information—including age, sex, diagnosis, and experimental group is collated in Table 1. For all epilepsy patients, the resected neocortical tissue used for recordings was located in the epileptic focus. For control patients, tissue was taken from the neocortex in order to gain access to deeper brain structures needing surgical treatment and displayed no structural/functional abnormalities. For our non-MCD cohort, 36 cells were analyzed from 12 patients, and for our MCD cohort, 44 cells were analyzed from 16 patients. We also analyzed 10 cells from 2 tumor control patients. In all cases, histopathological analyses confirmed diagnosis.

**Table 1:** Patient Information. Patient information regarding demographics, diagnosis, and resected brain area_categorized based on whether patient has malformation of cortical development (MCD). Acronyms: *APXA*, anaplastic pleomorphic xanthoastrocytoma, *FCD*, Focal Cortical Dysplasia, *DNET*, dysembryoplastic neuroepithelial tumor.

| Final diagnosis | MCD? | Resection location | Patient age at collection (years) | Patient sex |
| --- | --- | --- | --- | --- |
| APXA | Control | parietal lobe | 21 | M |
| Germinoma | Control | parietal lobe | 11 | M |
| FCD 1B | Yes | temporal tip | 9 | F |
| FCD IA | Yes | lateral temporal lobe | 17 | F |
| FCD IC | Yes | frontal lobe, temporal tip | 7 | M |
| FCD IIA | Yes | frontal lobe | 2 | F |
| Mild MCD | Yes | frontal lobe | 3 | F |
| Tuberous Sclerosis | Yes | frontal lobe | 10 | M |
| FCD IIA | Yes | occipital lobe | 19 | M |
| FCD IIB | Yes | frontal lobe | 2 | F |
| FCD IIID | Yes | lateral temporal lobe | 6 | M |
| FCD IIID | Yes | temporal lobe, frontal lobe | 18 | M |
| Tuberous Sclerosis | Yes | temporal lobe | 15 | M |
| FCD IIA | Yes | frontal lobe | 2 | F |
| FCD IIB | Yes | frontal lobe | 7 | F |
| Chaslin's subpial gliosis | No | lateral temporal lobe | 18 | M |
| Chaslin's subpial gliosis | No | lateral temporal lobe | 20 | M |
| Encephalomalacia | No | frontal lobe | 18 | M |
| Chaslin's subpial gliosis | No | lateral temporal lobe | 5 | F |
| Ganglioglioma grade I | No | temporal tip | 16 | F |
| Chaslin's subpial gliosis | No | frontal lobe | 5 | M |
| Chaslin's subpial gliosis | No | lateral temporal lobe | 10 | M |
| Gliosis and Meningioangiomatosis | No | parietal lobe | 16 | F |
| Gliosis | No | frontal lobe | 6 | M |
| Gliosis | No | temporal lobe | 5 | M |
| Low grade glial neoplasm | No | temporal lobe | 1 | F |
| DNET | No | temporal lobe | 10 | F |
| Glioma grade 1 (10/18/21) | No | Temporal lobe | 8 | M |
| Chaslin's subpial gliosis | No | temporal lobe, frontal lobe | 16 | M |
| DNET? | No | frontal lobe | 6 | F |

Based on previous research looking at *in vitro* ictal activity and bursting pyramidal neurons in L2/3 we decided to record from L2/3 for this study. L2/3 is thought to function as an integrator of information across cortical areas and play a role in regulating gain of cortical output (Luo et al., 2017, Quiquempoix et al., 2018)^2,40^. L2/3 retains a fairly homogenous class of pyramidal neurons, in contrast to deeper cortical layers (Mohan et al., 2015)^12^, although deep layer 3c neuron have been shown to have their own electrophysiologic and transcriptomic phenotypes (Moradi Chameh et al., 2021, Berg et al., 2020)^5,7^. We did not differentiate L2 from L3 neurons in this study.

### Diversity of L2/3 human epileptic neocortical pyramidal neuron morphology, action potential kinetics, and firing properties

L2/3 human pyramidal neurons are a diverse group of cells with various morphologies, electrophysiologic properties, and transcriptomic diversity making them difficult to characterize on electrophysiologic properties alone (Foehrig et al., 1991, DeFelipe et al., 1992, Deitcher et al.,2017, Kalmbach et al., 2018, Berg et al., 2020)^5,8,9,15,41^. In addition, the influence of epileptic seizures on neuron firing properties is not well understood (Levinson et al., 2020, Moradi Chameh et al., 2021)^6,7^.Therefore, in order to confirm that we were successfully targeting pyramidal cells in our epileptic tissue slices, a subset of neurons were filled with biocytin and stained with streptavidin conjugated to Alexa Fluor 593 or 488 and three-dimensional (3D) neuron reconstructions were performed. Figure 1 illustrates the diversity of L2/3 putative pyramidal neuron action potential (APs) shapes and firing patterns that were induced by depolarizing current steps (600ms). We found that L2/3 pyramidal neurons show diverse morphologies similar to those seen in normal human neocortical tissue (Mohan et al., 2015, Deitcher et al., 2017)^12,41^ indicating that gross morphology of epileptic L2/3 pyramidal neurons is intact (Fig. 1Aiii-Eiii). We also observed diverse spiking properties including differences in action potential shape including halfwidth, amplitude and AHP shape (Fig. 1Aii-Eii)and the propensity to fire continuously, accommodate quickly or stutter (Fig. 1Ai-Ei). This suggests that epileptic L2/3 pyramidal neuron morphology alone may not dictate neuron subtype and that the variability of epileptic L2/3 pyramidal neuron intrinsic properties may influence network excitability toward pro-ictal activity both *in vivo* and *in vitro*.

In order to differentiate between L2/3 pyramidal neuron subtypes of epileptic human tissue, we grouped neurons based on action potential shape or firing rate characteristics into 5 categories: accommodating neurons (n = 7, 4), regular spikers (RS) (n = 51, 21), stuttering neurons (n = 6, 4), early spikers (ES) (n = 9, 6), and “notch” neurons (n = 19, 10); named for their quick AHP as observed by Foehrig et al., 1991 and Lorenzon et al., 1992^8,10^. After our initial classification, we analyzed passive and active membrane properties (Figure 2) (Table 2) using a series of hyperpolarizing and depolarizing current steps (see Methods). We observed no significant differences between subtype passive membrane properties including resting membrane potential (RMP) (mV), AHP magnitude (mV), input resistance (MΩ) and voltage sag (%); however, we observed membrane decay (ms) differences between accommodating and RS neurons [KW test: p= 0.0468, H(4)= 9.650, Dunn test: accommodating vs RS (p= 0.0314, mean rank difference (MRD)= 31.49, z= 2.954)] (Fig. 2D-E, Table 2).

**Table 2.1:**
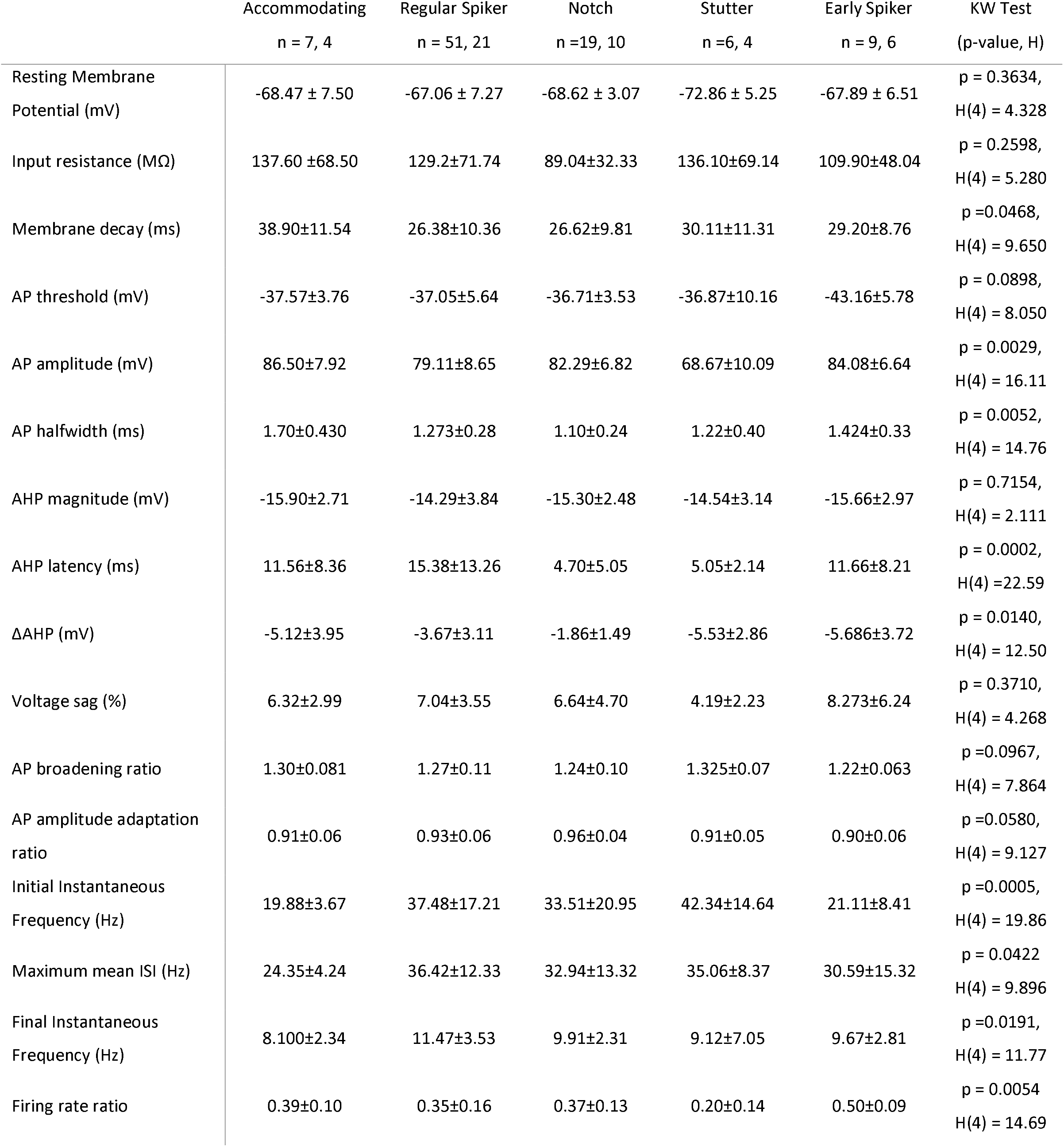
Differences in active and passive membrane properties among L2/3 neuron subtypes.

**Table 2.2:**
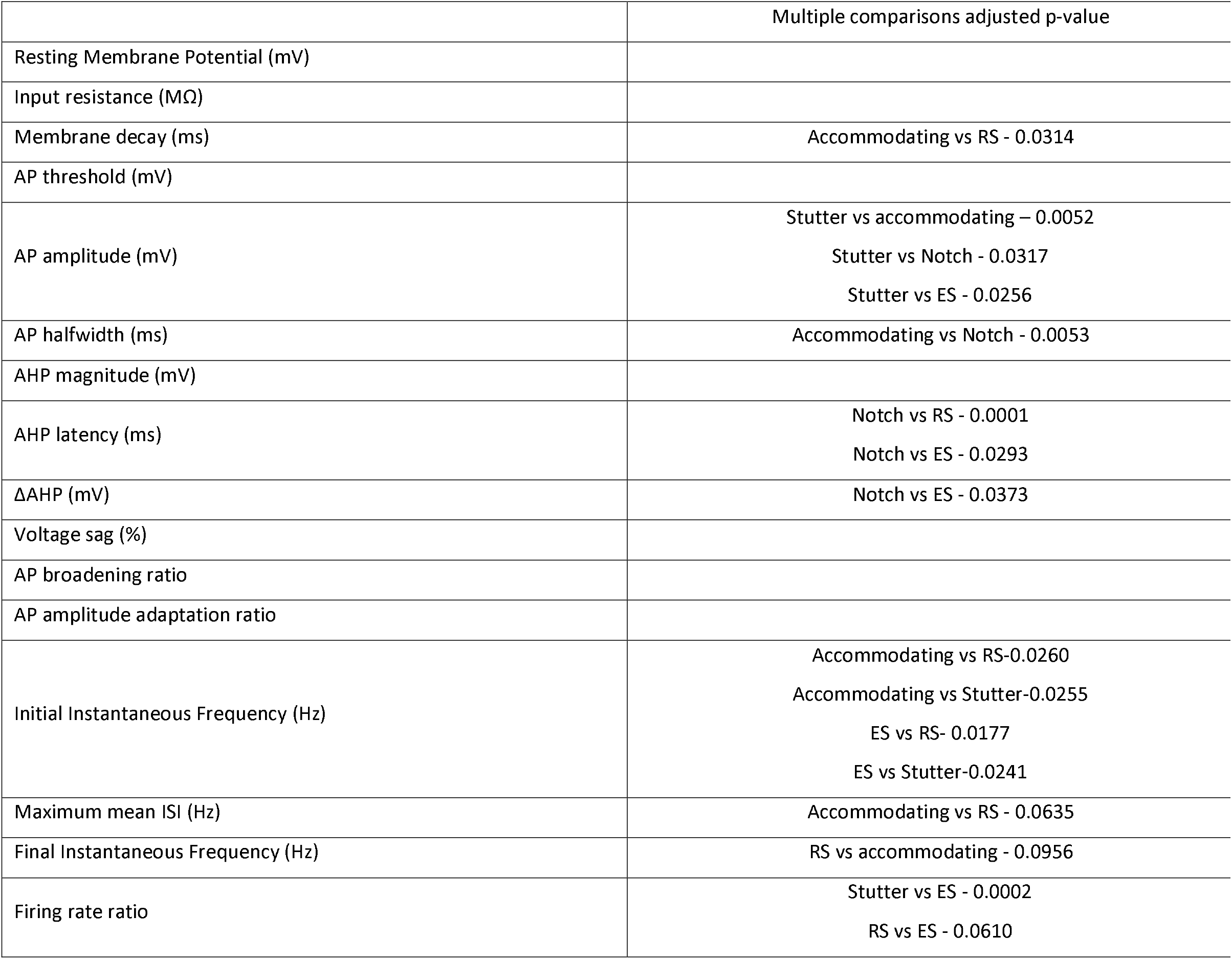
Post-hoc Dunn test p-values for statistically significant properties based on L2/3 neuron subtype. Intrinsic property data for neuron subtypes based on action potential shape and firing rate characteristics are presented as mean ± standard deviation (SD) (Table 2.1). Experimental numbers are reported as n= x, y where x is the number of neurons and y is the number of patients. The statistical comparison was a nonparametric Kruskal-Wallis test (KW) based on low n for some groups. Post-hoc Dunn test adjusted p-values are presented in Table 2.2.

We also observed differences in action potential properties between subtypes (Fig. 2A-C, E-N). Specifically, notch neurons showed the shortest halfwidth with a significant difference compared to accommodating neurons [KW test: p=0.0052, H(4) = 14.76, Dunn test: p=0.0053, MRD= 40.79, z= 3.464] (Fig. 2H), stutter neurons showed a reduction in AP spike amplitude (mV) with significant differences compared to accommodating, notch and ES neurons, respectively [KW test: p= 0.0029, H(4) = 16.11, Dunn test: stutter vs accommodating (p= 0.0052, MRD= 51.57, z= 3.472), stutter vs notch (p= 0.0317, MRD=3 6.89, z= 2.951), stutter vs ES (p= 0.0256, MRD= 42.44, z= 3.016)] (Fig. 2I), and notch neurons also had the quickest AHP latency (ms) with significant differences compared to RS and ES neurons [KW test: p= 0.0002, H(4) =22.59, Dunn test: notch vs RS (p= 0.0001, MRD= 31.90, z= 4.409), notch vs ES (p= 0.0293, MRD= 32.49, z= 2.974)] (Fig. 2F) and differences in ΔAHP compared to ES neurons (KW test: p=0.0140, H(4) = 12.50, Dunn test: p= 0.0373, MRD= 31.33, z= 2.900)] (Fig. 2G).

The action potential shape corresponded with differences between neuron subtype firing properties including initial instantaneous frequency [KW test: p= 0.0005, H(4) = 19.86] and FR ratio [KW test: p= 0.0054, H(4) = 14.69]. Interestingly, we observed that notch and stutter neurons trended toward quicker AHP latencies leading to a propensity to fire faster initially and accommodate their firing rate more rapidly (Fig 2J, Mi-Mii.). Regular spikers showed a diverse group of action potential shapes, but a large majority showed a similar trend toward faster AHPs lending the data set toward the same firing properties as notch and stutter neurons. Against our expectations, neuronal subtypes did not show any notable differences in max FR (Hz) or final instantaneous frequency (ms), and there was substantial overlap in action potential and firing rate property values between subtypes. Overall, these data suggest although these neurons may have differences in action potential and firing properties, they are likely part of the same pyramidal neuron subtype and not part of an interneuron subtype.

### AHP shape and latency doesn’t solely dictate L2/3 pyramidal neuron firing rate or firing rate accommodation

We then decided we wanted to differentiate these putative pyramidal neurons based solely on their AHP shape and latency, to make sure we weren’t biasing the subtypes based on our own characterization. AHPs reflect active ionic conductance after action potentials and dictates spike repolarization, instantaneous and repetitive firing, and FR accommodation and are controlled by different K= currents through channels such as BK, SK, and non-BK SK-mediated calcium-activated channels (Avoli et al., 1989, Lorenzon et al., 1992, Faber et al., 2002, Higgs et al., 2009, Rizzo et al., 2015, Mendez-Rodriguez et al., 2021)^4,10,42–44,45^. Previous research has shown that AHP shape in itself dictates firing properties with fAHP blockage leading to reduced FR accommodation (Avoli et al., 1991, Lorenzon et al., 1992, Foehrig et al., 1991) ^8,10,36^ and mAHP enhancement leading to increased initial instantaneous firing (Faber et al., 2002)^42^. In this study, AHP latency was determined based on the duration of time it took from threshold to the maximum AHP value. We found a subset of neurons had a fast AHP latency (fAHP) component that was ^~^5 ms or less and others only had a longer AHP latency value termed a medium AHP (mAHP) component.

We consequently wanted to determine if AHP shape was responsible for the marked differences in instantaneous firing and FR accommodation seen in neuronal subtype data (Fig. 2). To do this, we separated neurons based on AHP shape: those with fAHPs (n = 45, 20) and those with only mAHPs (n = 34, 14). We found that 39.24% of recorded neurons had fAHPs followed by a depolarization potential (ADP) and then by a mAHP (n = 31) (Fig. 3Ai) and 17.72% of neurons had fAHPs followed by a mAHP (n = 14) (Fig. 3Aii) as seen previously (Lorenzon et al., 1992 and Foehrig et al., 1991)^8,10^. 43.04% of recorded neurons were categorized as having mAHPs only (Fig. 3Aiii). Interestingly, L2/3 cortical pyramidal neurons in rats have been shown to have the fAHP followed by a fast ADP similar to Figure 3Ai that can trigger bursting which is a common phenotype seen in human epileptic brain tissue from cortex and hippocampus samples (Avoli et al., 1987, Higgs et al., 2009, Chen et al., 2011, Levinson et al., 2020)^6,32,43,46^.

We then compared active and passive properties between neurons with fAHPs to those without (mAHP neurons), collated in Figure 3 and Table 3, and found a significantly shorter AHP latency (Fig. 3D), as expected (MWU test: p < 0.000001, MD= 10.59, confidence interval(CI)= 9.050 to 13.60), and a more depolarizing ΔAHP in fAHP neuron compared to mAHP neurons (MWU test: p <0.0001,MD= −3.204, 95%CI= −4.608 to −2.289) (Fig. 3E). We also found fAHP neurons had a quicker max repolarizing slope (dV/dT) (unpaired t-test: p= 0.030, mean difference (MD)= 7.236 dV/dT, 95%CI= 0.7361 to 13.74) (Fig. 3Ci-Ciii) coordinating with shorter AP halfwidths (unpaired t-test: p= 0.006, MD= 0.208ms, 95%CI= 0.0615 to 0.354) (Fig. 3F) and sustained AP amplitude adaptation ratio (unpaired t-test: p= = 0.006, MD: −0.0322, 95%CI= −0.055 to −0.009) (Fig. 3G) indicating these fAHP neurons have a propensity for quicker action potentials that do not accommodate their spike amplitude during trains of APs compared to mAHP neurons. fAHP neurons had a propensity toward more depolarized ΔAHP with a subset reaching positive values previously described as a differentiating marker of LTS neurons in mouse layer 4 (Fig.3D) (Beierlein et al., 2003) ^47^. However, fAHP neurons also had lower hyperpolarization-induced sag (voltage sag ratio) compared to mAHP neurons (MWU: p= 0.0115, MD: 1.922, 95%CI= 0.5720 to 3.405) (Fig. 3H) and lower input resistance (MWU: p= 0.0323, MD= 50.17, 95%CI= 2.587 to 76.10) (Fig. 3I) indicating fAHP neurons are most likely not part of the LTS neuron subtype as these results contradict previous literature on regular LTS interneuron input resistance and voltage sag (Beierlein et al., 2003, Li et al., 2009, Tremblay et al., 2016, Levinson et al., 2020)^6,47–49^. The decrease in voltage sag of fAHP neurons may indicate deficits in HCN1 channels which play a role in Ih current, a contributor to enhanced frequency-dependent gain and possibly decreased excitability (Brennan et al., 2016, Herrmann et al., 2015, Kalmbach et al., 2018, Moradi Chameh et al., 2021)^7,15,50,51^.

**Table 3:** Intrinsic properties of neurons separated based on AHP shape and latency. Intrinsic properties of neuron subtypes split by AHP shape and latency are presented as mean ± standard deviation (SD). Experimental numbers are reported as n= x, y where x is the number of neurons and y is the number of patients. The statistical comparison was either an unpaired t-test or Mann Whitney U test (MWU) based on normality and variance.

|  | fAHP neurons<br>n = 45, 20 | mAHP neurons<br>n = 34, 14 | Statistical comparisons<br>(p-value) |
| --- | --- | --- | --- |
| Resting Membrane Potential (mV) | -68.68±6.36 | -66.05±6.71 | MWU<br>p = 0.0492 |
| Input resistance (MΩ) | 103.40±41.80 | 150.40±81.61 | MWU<br>p = 0.0323 |
| Membrane decay (ms) | 27.23±8.65 | 27.54±7.91 | Unpaired t-test<br>p = 0.8712 |
| AP threshold (mV) | -36.43±5.45 | -37.70±5.54 | MWU<br>p = 0.5612 |
| AP amplitude (mV) | 83.22±8.02 | 84.48±9.56 | MWU<br>p = 0.5965 |
| AP halfwidth (ms) | 1.21±0.28 | 1.422±0.35 | Unpaired t-test<br>p = 0.0060 |
| AHP magnitude (mV) | -15.29±2.97 | -14.93±3.40 | MWU<br>p = 0.6197 |
| AHP latency (ms) | 3.91±1.08 | 18.75±12.31 | MWU<br>p<0.000001 |
| ΔAHP (mV) | -2.00±2.21 | -5.45±2.82 | MWU<br>p = <0.0001 |
| Voltage sag (%) | 5.68±4.13 | 7.17±2.97 | MWU<br>p = 0.0115 |
| AP broadening ratio | 1.26±0.11 | 1.27±0.098 | MWU<br>p = 0.4286 |
| AP amplitude adaptation ratio | 0.94±0.04 | 0.91±0.060 | Unpaired t-test<br>p = 0.0064 |
| Initial Instantaneous Frequency (Hz) | 32.51±18.31 | 30.71±12.57 | MWU<br>p =0.9011 |
| Maximum firingrate (Hz) | 27.61±6.78 | 28.15±9.48 | MWU<br>p = 0.9438 |
| Final Instantaneous Frequency (Hz) | 9.99±2.69 | 11.53±4.50 | MWU<br>p = 0.1650 |
| Firing rate ratio | 0.39±0.15 | 0.40±0.13 | Unpaired t-test<br>p = 0.6957 |

Against our expectations, we saw no significant differences in initial or final instantaneous firing, Max FR, or FR accommodation; although there were trends toward an elevated F/I curve for both mean and final frequency of mAHP neurons over the depolarizing current steps. (Fig.3I-L, Table 3). These findings are opposite those previously observed and negates our hypothesis and overarching theory that the AHP shape and latency, and the underlying channels that dictate this phenotype completely controls human neocortical L2/3 pyramidal neuron firing rate and adaptation significantly (Avoli et al., 1989, Lorenzon et al., 2002, Foehrig et al. 1991). We postulate that it is a combination of factors including voltage-gated sodium and potassium channel kinetics and expression and passive properties including input resistance, membrane capacitance and Ih current (Howell et al., 2015, Ceballos et al., 2016)^52,53^. It is worth noting that with almost 40% of these pyramidal neurons having ADPs, which can help neurons enter bursting mode, it would be worthwhile to interrogate if these neurons are more likely to become the paroxysmal discharging neurons we see in epileptic human tissue since they are thought to play a pivotal role in generating ictal activity *in vitro* and *in vivo* (Levinson et al., 2020).

### Epileptic subtype influences L2/3 pyramidal neuron intrinsic properties compared to tumor control neocortical tissue

The etiology of epilepsy is immense and has been studied in vitro for the past few decades (Abdijadid et al., 2015, Avoli et al, 1991a, 1991b, 1994, Calcagnotto et al., 2005, Chang et al., 2019, Chen et al. 2011, Cossart et al., 2001, Heinemann et al., 2014, Iffland and Crino, 2017, Jozwiak et al., 2006, Lasarge et al., 2014, Le Duiguo et al., 2018, Levinson et al., 2020, Lipton et al., 2014, Nakagawa et al., 2017, Lipton et al., 2014, O’Dell et al., 2012, Hodge et al., 2019, Saxena et al., 2015, Schwartzkroin et al., 1994, Schwartkroin et al., 1983, Schwarz et al., 2019, Ting et al., 2018a, 2018b, Toth et al., 2018, Yasin et al., 2013) ^3,6,21,24–26,28,30,36,38,46,54–69^. With each etiology leading to differences in neuronal migration, differentiation, and synaptic development, a wide array of neuronal intrinsic properties and network level properties emerge. In order to begin to differentiate between epileptic subtypes, we next categorized L2/3 pyramidal neurons based on whether the patient’s epilepsy was a malformation of cortical development (MCD) or not (non-MCD). We did not further differentiate neurons based on morphology or physiology in order to get a general picture of how L2/3 pyramidal neurons differed between epileptic subtypes.

In order to characterize L2/3 pyramidal neurons from MCD, non-MCD and control acute slices, we ran a depolarizing current ramp protocol to assess resting membrane potential (mV) (RMP) and AP rheobase (mV) and depolarizing and hyperpolarizing current steps to determine active and passive membrane properties (Fig. 4A-B). We found that L2/3 pyramidal neurons from both MCD and non-MCD epileptic tissue showed substantial differences in passive membrane properties compared to control and varying differences in action potential properties compared to control and each other (Fig. 4, Table 4). Compared to control, we observed an increase in membrane decay [KW test: p= = 0.0216, H(2) = 7.674, Dunn test: (control vs MCD, p= 0.0059, MRD= −22.79, z= 2.407), (control vs non-MCD, p= 0.0177, MRD= −25.45, z= 2.754)] (Fig. 4E), a decrease in voltage sag ratio [KW test: p= 0.0111, H(2)= 8.994, Dunn test: (control vs MCD, p= 0.0115, MRD= 27.82, z= 2.892), (control vs non-MCD, p= 0.0154, MRD= 26.35, z= 2.798)] (Fig.4F) and no difference in input resistance compared to control [KW test: p= 0.0287, H(2)= 7.101, Dunn test: (control vs MCD, p= 0.5066, MRD= −13.24, z= 1.376), (control vs non-MCD, p> 0.9999, MRD= 2.357, z= 0.2502)] (Fig. 4D) for both MCD and non-MCD neurons, indicating it takes longer durations for current to elicit voltage changes (membrane decay) and possible aberration in HCN1 channels (voltage sag ratio), respectively. These epileptic pyramidal neuron property differences could be due to the Ih current partially determined by HCN1 channel activity and/or aberrations in voltagegated channel kinetic.

**Table 4.1:**
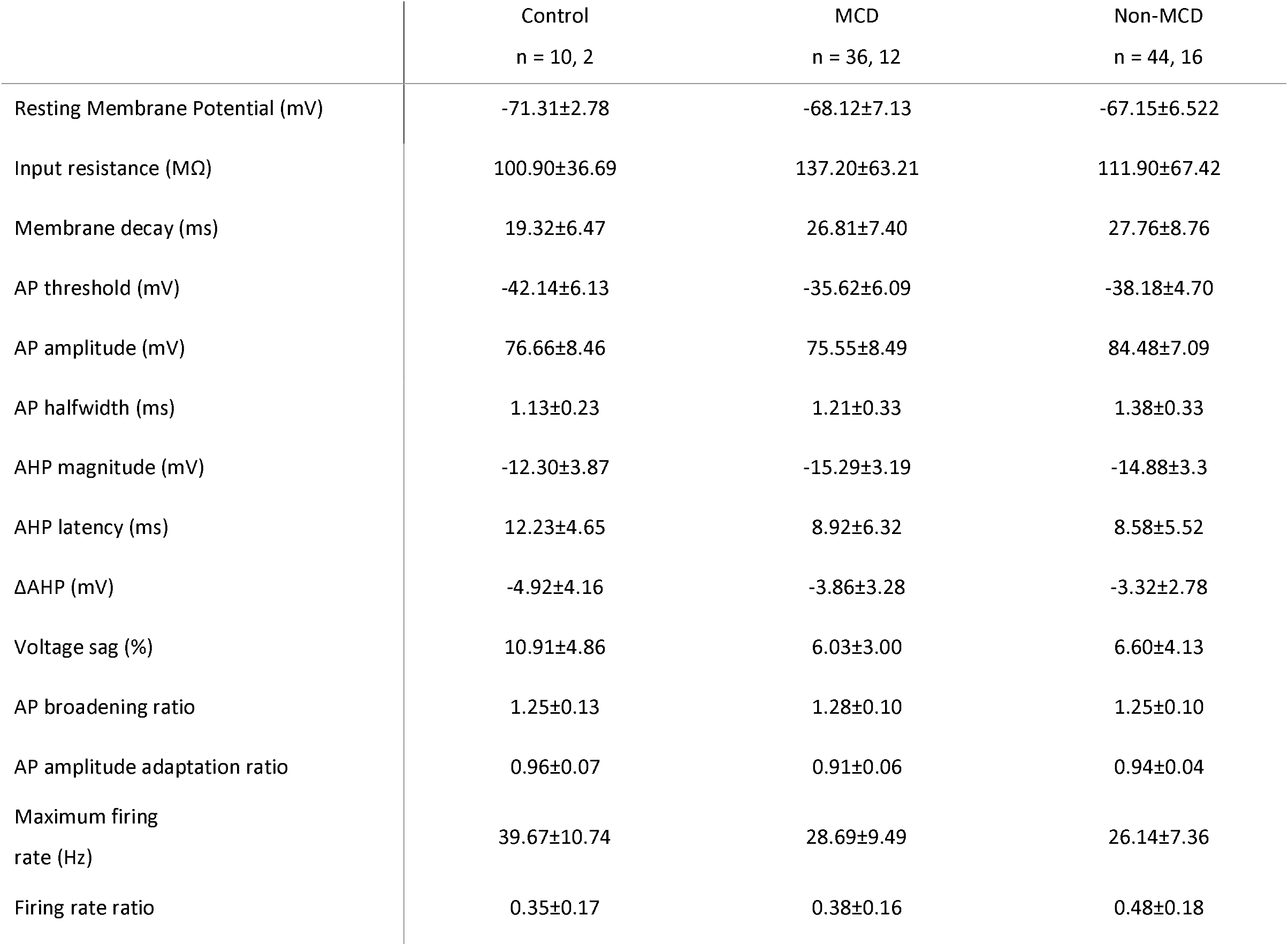
Intrinsic properties of L2/3 pyramidal neurons split by control and epilepsy subtype.

**Table 4.2:**
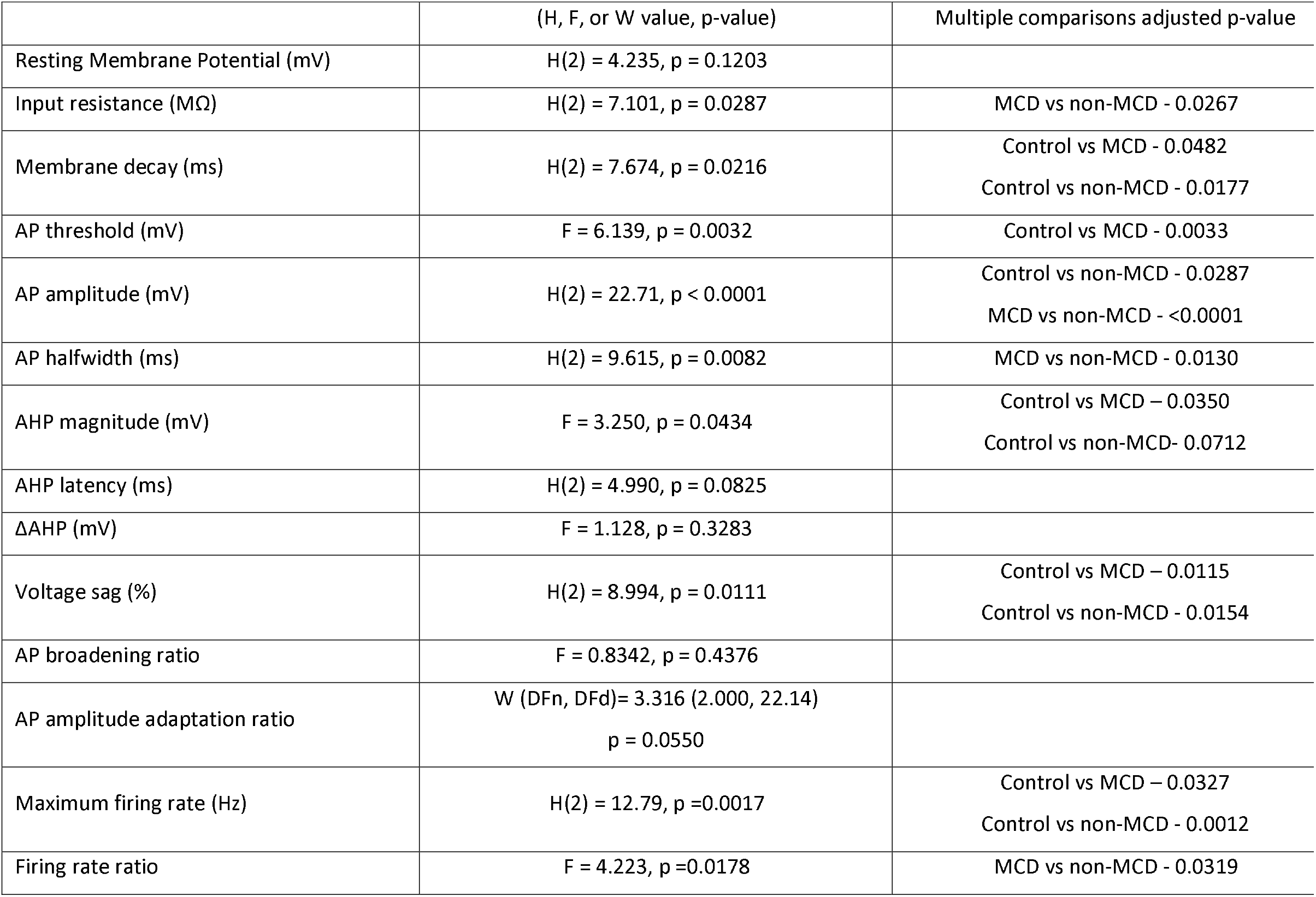
Group comparison values with post hoc adjusted p-values for statistically significant properties based on epilepsy subtype. Intrinsic properties based on control vs epileptic subtype are presented as mean ± standard deviation (SD). Experimental numbers are reported as n= x, y where x is the number of neurons and y is the number of patients. The statistical comparison was either an ANOVA, Kruskal-Wallis test (KW) or Welch’s ANOVA based on normality and variance (Table 4.1). Post hoc KW H values, ANOVA F values or Welch’s W values are presented along with correct adjusted p-values are presented in Table 4.2.

Overall, these property deficits compared to control would cause L2/3 neurons to have improper subthreshold oscillations and lack of temporally precise input computation leading to deficits in network synchronization and input summation which can lead to overall hyperexcitability (Higgs et al., 2009, Ceballos et al., 2016, Moradi Chameh et al., 2021) ^7,43,52^. Furthermore, several neurons, including those in both epileptic subtypes, have longer halfwidths (ms) [KW test: p= 0.0082, Dunn test: (control vs MCD, p= 0.8741), (control vs non-MCD, p= 0.0491)] (Fig. 4L) compared to control neurons. We indicate no differences in a number of other action potential and firing rate adaptation properties compared to control, including: AHP latency, ΔAHP or AP amplitude adaptation and FR ratio between control neurons and those from epileptic foci (Table 4). However, the differing action potential properties and reduction in max FR indicate that L2/3 epileptic pyramidal neurons show signs of dysregulated computation compared to control neurons.

We then looked at various action potential and firing properties and found further indication of perturbations of L2/3 pyramidal neurons from MCD and non-MCD compared to control. Specifically, compared to control both epileptic subtypes had decreased max firing rates (Hz) [KW: p= = 0.0017, H(2) = 12.79, Dunn test: (control vs MCD, p= 0.0327, MRD= 23.68, z= 2.546), (control vs non-MCD, p= 0.0012, MRD= 32.47, z= 3.544)] (Fig. 4M, right). This indicates that when concatenated, MCD and non-MCD L2/3 pyramidal neurons show signs of being hypoexcitable compared to control. This may be due to intrinsic developmental problems caused by a homeostatic changes due to hyperexcitability across the network or due to improper neuron migration, development, and synaptic plasticity all of which have been shown to influence neuron excitability (Medvedeva et al., 2020, Jarero-Basulto et al., 2018, Park et al., 2018, Curatolo et al., 2018, Andreae and Burrone, 2014)^20,22,70–72^. Moreover, this computational deficit phenotype across epilepsy subtypes may be due to an overabundance of excitatory bursting neurons that facilitate network synchronization and are thought to induce ictal activity in both MCD and non-MCD epilepsies (Levinson et al., 2020). To add to this, L2/3 MCD neurons have a significantly more depolarized threshold (mV) [ANOVA: p= 0.0032, F= 6.139, Tukey HSD: control vs MCD, p= 0.0033, MD= −6.520mV, 95%CI= −11.15 to −1.889)] (Fig. 4I) and an increase in AHP magnitude (mV) compared to control [ANOVA: p= 0.0434, F= 3.250, Tukey HSD: (control vs MCD, p= 0.0350, MD= 2.985mV, 95%CI= 0.1702 to 5.800)] (Fig. 4J) indicating more hyperpolarizing potentials post AP, meaning more need for depolarizing potentials to spike continuously.

### Epilepsy etiology influences L2/3 pyramidal neuron properties

In non-MCD epilepsies, we observed substantial increases in halfwidth [KW test: p= 0.0082, H(2) = 9.615, Dunn test: (MCD vs non-MCD, p= 0.0130, MRD= −16.97, z= 2.854)] and lack of FR accommodation compared to MCD epilepsy [ANOVA: p= 0.0178, F = 4.223, Tukey: (MCD vs non-MCD, p= 0.0319, MD= − 0.09999, 95%CI= −0.1929 to −0.007073)], with trends toward less FR accommodation compared to control (Fig. 4N), and an increase in AP amplitude compared to both control and the MCD subtype [KW test: p< 0.0001, H(2) = 22.71, Dunn test: (control vs non-MCD, p= 0.0287, MRD= −24.14, z= 2.591), MCD vs non-MCD, p <0.0001, MRD= −27.13, z= 4.566)]. This indicates to us that non-MCD L2/3 pyramidal neurons do show indications of differences with MCD neurons in terms of their firing properties, i.e., they fire just as frequently (still less than control pyramidal neurons) yet sustain this firing with longer halfwidths and larger AP amplitudes than MCD L2/3 pyramidal neurons. This could allow more neurotransmitter release and could work in concert with bursting neurons and the improper proportion of interneuron subtypes observed in MCDs, such as FCD (D’Antuono et al., 2004, Liang et al., 2020)^19^, as improper synchrony mixed with larger and longer action potentials that generate excitatory neurotransmission could help generate more substantial network excitability (Yang et al., 2006)^73^.

We postulate that this may be due to somatic mutations that cause migration issues and network maturation deficiencies leading to cell differentiation deficits (Guerrini et al., 2015, Brennan et al., 2016, Subramanian et al., 2020, Iffland and Crino, 2017)^29,30,50,74^. In addition, MCD encompasses FCD and TSC which both have been characterized as having cytomegalic dysmorphic neurons, immature pyramidal neurons, and balloon/giant neurons with a variety of input resistances and capacitances which together influence the MCD L2/3 neurons intrinsic properties (Levinson et al., 2020, Crino, 2015)^6,27^.

### 4-AP influences L2/3 pyramidal neuron intrinsic properties

4-aminopyridine (4-AP) is a convulsant drug that blocks A-type K+ current, thereby facilitating neurotransmitter release at both excitatory and inhibitory synapses and has been shown to block fAHPs and increase AP halfwidth to dictate neuron firing rate and firing adaptation (Lorenzon et al., 1992, Faber et al., 2002, Guerrini et al., 2015)^10,42,29^. 4-AP has also been used in concert with a mg2+ free aCSF, high extracellular K+, GABA, glutamate, and inhibitory neurotransmitter blockers to promote and induce interictal and ictal activity in human epilepsy brain slice studies (Avoli et al.,1991, Avoli et al., 1998, Cepeda et al., 2018, Chang et al., 2019, Guerrini et al., 2015 Levinson et al., 2020) ^10,17,46,33,54,55^. However, to our knowledge, no substantial work has gone into how 4-AP influences L2/3 pyramidal neurons from epileptic acute slices. Since 4-AP is so widely used in *in vitro* epilepsy electrophysiology recordings, we wanted to characterize the intrinsic properties it influences to understand how it can lead to proepileptic conditions.

To this end, we performed whole-cell current clamp recordings on L2/3 pyramidal neurons and elicited hyperpolarizing and depolarizing current steps before and after 4-AP (100μM) wash on. When comparing neurons before and after 4-AP wash on (n=17, 8), we found that 4-AP induced interesting alterations in L2/3 pyramidal neuron passive membrane properties, action potential properties and firing properties (Fig. 5, Table 5). As expected, it produced an increase in halfwidth (paired t-test: p= 0.0091, MD= 0.3227ms, 95%CI= 0.09211 to 0.5534) (Fig. 5A-B,D) and an increase in AHP latency (paired t-test: p= 0.0002, MD= 6.940ms, 95%CI= 3.846 to 10.03) (Fig. 5G) corresponding with a decrease in AHP magnitude (Wilcoxon test: p= 0.0021, median of difference= 2.164, 95%CI= 1.122 to 3.682) (Fig. 5F) indicating 4-AP was blocking fAHPs and influencing voltage-gated K+ channels.

**Table 5:**
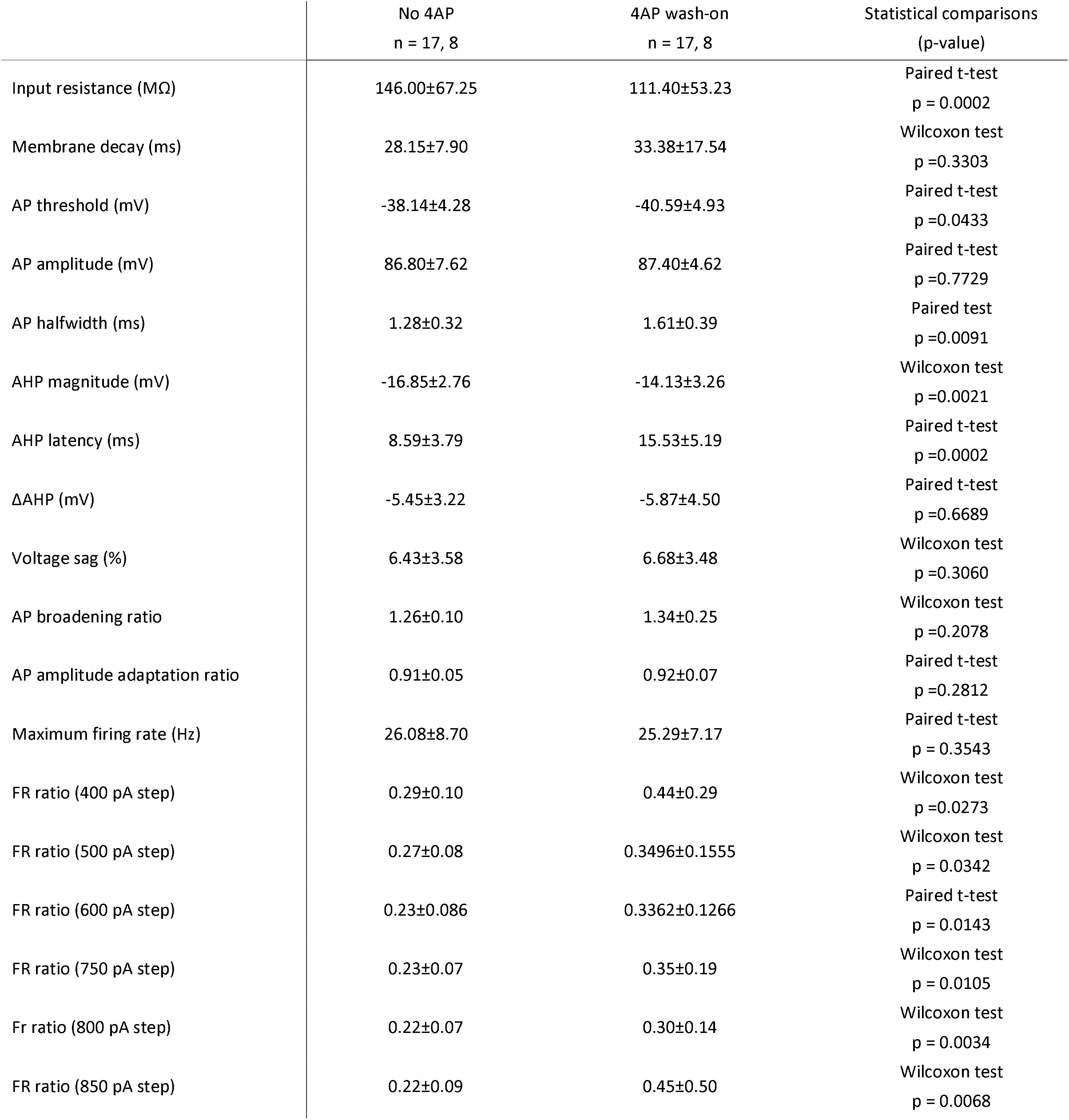
4AP induces changes of L2/3 pyramidal neuron intrinsic properties. Data are presented as mean ± standard deviation (SD). Experimental numbers are reported as n= x, y where x is the number of neurons and y is the number of patients. The statistical comparison was either a paired t-test for normally distributed data or a Wilcoxon matched-pairs signed rank test for non-normal data or if the data was normal but SD differed by more than 2X.

4-AP also induced a significant decrease in L2/3 neuron input resistance (paired t-test: p= 0.0002, MD= −34.64ms, 95%CI= −50.10 to −19.19 MΩ) (Fig. 5C) and a shift to a more negative threshold for AP generation (paired t-test: p= 0.0433, MD=-2.450mV, 95%CI= −4.816 to −0.08288) (Fig. 5E). 4-AP had no significant effect on voltage sag (Wilcoxon test: p=0.3060) (Table 5) or AP amplitude (paired t-test: p= 0.7729) (Fig. 5H) or max FR (paired t-test: p= 0.3543) (Fig. 5I). However, it did cause an increase in EPSP frequency (not shown) and caused significant fluctuation of FR accommodation at more depolarizing potentials (400, 500, 600, 750, 800, 850pA steps) due to approximately 1 second long sporadic hyperpolarizations [(400pA step: Wilcoxon test: p= 0.0273, median of differences: 0.05926, 95%CI= 0.0007553 to 0.3011), (500pA step: Wilcoxon test: p 0.0342, median of differences= 0.008756, 95%CI= 0.002787 to 0.1029), (600pA step: paired t-test: p= 0.0143, MD= 0.09877, 95%CI= 0.02299 to 0.1745), (750pA step: Wilcoxon test: p= 0.0105, median of difference= 0.03876, 95%CI= 0.02685 to 0.2142), (800pA step: Wilcoxon test: p= 0.0034, median of difference= 0.03472, 95%CI= 0.02125 to 0.1535), (850pA step: Wilcoxon test: p= 0.0068, median of difference= 0.02957, 95%CI= −0.05532 to 0.5216)] (Fig. 5J).

Overall, 4-AP causes L2/3 pyramidal neurons to shift to spiking at more hyperpolarized potentials and leads to sporadic repetitive firing yet has no influence on L2/3 pyramidal neuron max FR (Hz). With the increase in halfwidth (paired t-test: p= 0.0091, MD= 0.3227 ms, 95%CI= 0.09211 to 0.5534) and no significant change of AP amplitude, 4-AP allows L2/3 pyramidal neurons to fire longer action potentials at the same efficiency most likely leading to an increase in glutamatergic neurotransmission (Yang et al., 2006)^73^. Furthermore, the sporadic FR accommodation allows these neurons to be turned on and off and when switching to “on” mode they recuperate to normal firing rates. Whether or not these cells turn on and off together has not yet been assessed. Overall, this follows a similar trend to Figure 4 yet more drastically exacerbates a cortical environment where properties influencing input summation and synchronization such as firing rate accommodation and action potential halfwidth may work in concert to elicit more glutamatergic neurotransmission.

## Discussion

In the present study, we used cortical tissue from pediatric epilepsy surgery patients to characterize L2/3 pyramidal neurons located within the epileptic foci more conclusively. Most previous work in L2/3 has included other layers of interest within the analysis, only characterized rudimentary properties, or recorded from pyramidal neurons outside of the epileptic foci and concluded they were from “normal” human cortical tissue. In this study, we also wanted to determine if malformation of cortical development (MCD) epilepsies had significant differences in intrinsic properties compared to non-MCD epilepsies based on differences in underlying cellular mechanisms caused by specific genetic perturbations and differences in neuronal developmental and maturation.

To our knowledge, this is the first comprehensive study focused on characterizing electrophysiological and morphological properties of human L2/3 neocortical neuron from within the epileptic focus and comparing these neurons to normal brain tissue taken from patients without any history of epilepsy. We determined that L2/3 pyramidal neurons show a fast variety of spiking patterns and AP kinetics (Fig. 1). We initially believed the notch neuron subtype to be similar to LTS interneurons, which usually express cholecystokinin (CCK), calretinin (CR) or somatostatin (SST) and have a fAHP followed by an ADP(Beierlein et al., 2003, Tremblay et al., 2016)^49^. We were particularly interested in these because of the AHP shape looking similar to LTS interneurons, but also because it was originally viewed under DIC as pyramidal. Could there be an issue with AP maturation or neuron differentiation and migration within the epileptic foci? For it has already been found that interneuron population levels are altered in subtypes of FCD, specifically a reduction in PV-interneurons (Liang et al., 2020). Interestingly, however, after 3D reconstruction of notch neuron morphology and analysis of their active and passive membrane properties we found these were pyramidal neurons and may be a class of L2/3 pyramidal neurons not really observed in mouse cortex (Fig. 2).

We also noted from past research, a dedication to AHP latency and shape as a dictator of firing rate and firing adaptation. Interestingly, compared to previous reports in human L2/3 looking at the channels that dictate AHP and how SK, BK, and other calcium-activated K+ channels regulate AHPs and therefore, neuron firing patterns Avoli et al., 1989, Foehring et al., 1991, Lorenzon et al., 1992, Faber et al., 2002, Higgs et al., 2009, Rizzo et al., 2015, Mendez-Rodriguez et al., 2021), we found neurons with fAHPs (^~^5ms to max AHP compared to time of threshold (ms)) compared to those with longer AHP latencies and no ADP (mAHP neurons) indicating having certain AHP components does not completely dictate firing properties of L2/3 pyramidal neurons, but may be a good way to differentiate human L2/3 pyramidal neurons going forward based on variety of AP properties and passive membrane properties that differed between these two cell types (Fig. 3). Future research will hopefully indicate if these L2/3 neurons are their own subtype with specific physiological roles in the hyper-synchronized network and if these neurons may be the pyramidal neurons that are more likely to become bursters because of their unique AHP components and ADPs which have been shown to help neurons transition into bursters (Higgs et al., 2009, Chen et al., 2011)^43,46^.

Furthermore, intrinsic properties have been reported comparing between subtypes of MCD and non-MCD groups (Cepeda et al., 2012, Cepeda et al., 2018)^6,16,17^ with no significant differences in intrinsic properties previously stated between pyramid neurons. However, whole-cell patch clamp recordings were obtained from layers outside of L2/3 and we know now that L2/3 vs deeper layers differ in intrinsic excitability and membrane properties (Moradi Chameh et al., 2021). Therefore, this study is the first to fully characterize the electrophysiological differences between L2/3 pyramidal neurons from control tissue and MCD and non-MCD epileptic focal tissue (Fig. 4). Intriguingly, we found a significant increase in membrane decay (ms), decrease in voltage sag (%) and decrease in max FR rate (Hz) between control L2/3 pyramidal neurons and those within the MCD and non-MCD subtype. Indicating that it takes more current to elicit the same voltage response in these epileptic pyramidal neurons, that HCN1-mediated current (Ih current) is aberrant leading to subthreshold oscillation deficits in the epileptic pyramidal neurons and a decrease in max FR may indicate a homeostatic mechanism in which the pyramidal neurons fire less frequently, but each AP has a longer duration leading to a compensatory increase in pyramidal neuron excitability.

When comparing MCD neurons with control, we also found a shift to more depolarized threshold and larger AHP. Interestingly, Powell et al., 2021 illustrated recently that with an increase in extracellular concentration of [K+], AHP magnitude got bigger and threshold got slightly more depolarized ^75^. It could therefore be there is more outflux of [K+] into the extracellular space because of the hyperexcitable cortical slice compared to control (Avoli et al., 2005). Furthermore, this trend also is the case with non-MCD slices as they show signs of an increase in AHP magnitude and shift to a more depolarized threshold (mV).

Non-MCD L2/3 pyramidal neurons also showed a significant increase in AP amplitude compared to control and increased AP amplitude, AP halfwidth, and less FR accommodation compared to MCD L2/3 neurons (Fig. 4). Overall, these differences are subtle between epilepsy subtypes and show correlation with each other in terms of their intrinsic properties compared to control neurons. Overall, although there are slight differences, we see that these epileptic L2/3 neurons fire less often, have longer AP durations, don’t accommodate their FR as much, and have issues with membrane properties including ability to fire the same when given the same amount of current as control neurons (membrane decay) and having aberration in their HCN1 channels, which has been shown to cause enhanced excitability in L2/3 pyramid neurons in rats (Strauss et al., 2004)^76^.

Lastly, the various influences 4-AP has on neuron AP AHPs and its different influences on mouse excitatory and inhibitory neurons (Williams and Hablitz. 2015)^34^ is crucial in order to understand not just that we can use 4-AP in order to induce interictal and ictal activity in human cortical slices, but to understand how it effects the underlying neurons. As expected, we observed that 4-AP causes an increase in AP halfwidth, increase in AHP latency, and decrease in AHP magnitude. Indicating again, longer duration APs with blockage of [K+]. In addition, we noted a decrease in input resistance (Fig. 5C) which has not been previously reported and interestingly no change in max FR only an increase and fluctuation of FR accommodation (Fig. 5J). This was due to 4-AP causing hyperpolarizing oscillations during the current steps and therefore allowed these neurons to turn on and off, with on periods being a recouped neuron that had the ability to continue sustained firing instead of going through voltagegated Na+ channel rundown.

Overall, we saw a trend and expedited phenotype when comparing epileptic subtype to control (Fig. 4) and neuron intrinsic properties before and after 4-AP wash on (Fig. 5). AP halfwidth duration gets significantly longer, allowing for more glutamatergic neurotransmission (Yang et al., 2006), [K+] in the extracellular space is most likely higher (Avoli et al., 2005) or K+ channels are blocked so we observe changes in AHP latency and magnitude, and lastly, FR accommodation happens less so compared to controls. This is because of the reduction in max firing overall or the ability for the neuron to fluctuate between on and off with the 4-AP oscillations. Together, these data indicate that L2/3 pyramidal neurons from human epileptic foci are overall don’t have proper temporal summation and sustain themselves easily during higher doses of excitation and therefore, may release more glutamate into the epileptic-like *in vitro* network. Future research should indicate if AP halfwidth and glutamate release probability is tied in human pyramidal neocortical neurons. In addition, research looking at how interneurons are influenced by 4-AP and how different interneuron subtypes are dictating the shift between ictal and interictal states in slice is needed deeply.

